# Direct observation of coordinated assembly of individual native centromeric nucleosomes

**DOI:** 10.1101/2023.01.20.524981

**Authors:** Andrew R. Popchock, Joshua D. Larson, Julien Dubrulle, Charles L. Asbury, Sue Biggins

## Abstract

Eukaryotic chromosome segregation requires the kinetochore, a megadalton-sized machine that forms on specialized centromeric chromatin containing CENP-A, a histone H3 variant. CENP-A deposition requires a chaperone protein HJURP that targets it to the centromere, but it has remained unclear whether HJURP has additional functions beyond CENP-A targeting and why high AT DNA content, which disfavors nucleosome assembly, is widely conserved at centromeres. To overcome the difficulties of studying nucleosome formation in vivo, we developed a microscopy assay that enables direct observation of de novo centromeric nucleosome recruitment and maintenance with single molecule resolution. Using this assay, we discover that CENP-A can arrive at centromeres without its dedicated centromere-specific chaperone HJURP, but stable incorporation depends on HJURP and additional DNA-binding proteins of the inner kinetochore. We also show that homopolymer AT runs in the yeast centromeres are essential for efficient CENP-A deposition. Together, our findings reveal requirements for stable nucleosome formation and provide a foundation for further studies of the assembly and dynamics of native kinetochore complexes.

## Introduction

Replicated chromosomes must be accurately segregated to opposite poles during mitosis, a process that relies on their attachment to mitotic spindle microtubules via a conserved megadalton-sized protein network called the kinetochore (Biggins, 2013; Cheeseman, 2014; Kixmoeller *et al*, 2020; Musacchio & Desai, 2017; Santaguida & Musacchio, 2009). Errors in this process can lead to the rapid accumulation of mis segregated chromosomes resulting in a cellular condition called aneuploidy, a hallmark of cancerous cells (Gordon *et al*, 2012; Herman *et al*, 2015; Holland & Cleveland, 2009; Sheltzer *et al*, 2011). To ensure the fidelity of this process, kinetochores are assembled each cell cycle onto defined regions of chromosomes called centromeres (Clarke & Carbon, 1985; Cleveland *et al*, 2003; McAinsh & Marston, 2022). Among different organisms, centromeres vary in size and architecture and are epigenetically defined by the recruitment of a specialized H3 histone variant called CENP-A (Palmer *et al*, 1991; Sullivan *et al*, 1994). The centromere-specific targeting and deposition of CENP-A relies upon an essential conserved chaperone protein, HJURP (Camahort *et al*, 2007; Dunleavy *et al*, 2009; Foltz *et al*, 2009; Mizuguchi *et al*, 2007; Stoler *et al*, 2007). Once established, this specialized CENP-A nucleosome is then recognized by specific kinetochore proteins that enable complete kinetochore complex formation (McKinley & Cheeseman, 2016). CENP-A deposition onto centromeres is tightly regulated in cells as ectopic mis-incorporation contributes to chromosomal instability (CIN) (Shrestha *et al*, 2017). Consistent with this, CENP-A and HJURP overexpression, which can be driven by p53 loss, are common among various cancer types and have emerged as therapeutic cancer targets because higher levels are correlated with poor prognosis (Filipescu *et al*, 2017; Mahlke & Nechemia-Arbely, 2020).

Centromeric nucleosomes are critical for chromosome segregation, so it is surprising that the only widely conserved feature of centromeric DNA, its AT-rich content, is canonically a poor template for nucleosome assembly in reconstitutions with purified recombinant proteins (Kunkel & Martinson, 1981; Prunell, 1982). As a result of this, reconstitutions of assembled centromeric histones with centromeric DNA *in vitro* results in intrinsically unstable nucleosomes, making it difficult to study the functional role of the AT-rich centromeric DNA (Dechassa *et al*, 2011; Dechassa *et al*, 2014; Xiao *et al*, 2011). Recent breakthrough structural studies of CENP-A nucleosomes assembled with native centromeric DNA have found a more loosely associated centromeric DNA-nucleosome complex, which may provide distinct binding sites for the recruitment of DNA-binding kinetochore proteins (Guan *et al*, 2021; Zhou *et al*, 2019). However, these reconstitutions required the use of a single-chain antibody fragment (scFv) to stabilize the nucleosome in both yeast and human reconstitutions (Guan *et al*., 2021; Zhou *et al*., 2019). More complex reconstitutions that included additional kinetochore proteins required modification of the centromeric DNA sequence to include the nucleosome positioning Widom 601 DNA (Dendooven *et al*, 2022; Guan *et al*., 2021; Lowary & Widom, 1998; Yan *et al*, 2018), underscoring the difficulty of reconstituting stable kinetochore structures on centromeric DNA. While a more complete reconstitution of the constitutively centromere associated network (CCAN) kinetochore proteins assembled onto a CENP-A nucleosome was achieved on α-satellite DNA, the functional role of AT-rich centromeric DNA remains unclear (Yatskevich *et al*, 2022).

In contrast to most eukaryotes, budding yeast have sequence-specific point centromeres consisting of similar but not identical ∼125 bp DNA segments containing three different <u>c</u>entromere-<u>d</u>efining <u>e</u>lements, CDEI, CDEII and CDEIII (Biggins, 2013; Carbon, 1984; Carbon & Clarke, 1984). CDEI and CDEIII have consensus sites to recruit the centromere binding factors 1 and 3 (Cbf1 and CBF3), respectively. One function of Cbf1 is to protect centromeres from transcription to ensure chromosome stability (Cai & Davis, 1990; Hedouin *et al*, 2022), while the CBF3 complex (consisting of Ctf3, Cep3, Skp1 and Ndc10) coordinates with the Cse4 (CENP-A in humans) specific chaperone Scm3 (HJURP in humans) to promote the deposition of Cse4^CENP-A^ at CDEII (Camahort *et al*., 2007; Cho & Harrison, 2011b; Cole *et al*, 2011; Guan *et al*., 2021). Similar to other organisms, the CDEII element lacks sequence homology but consists of highly AT-rich DNA (Guan *et al*., 2021). Changes in the length or AT-content of CDEII compromise centromere stability *in vivo* for unknown reasons (Clarke & Carbon, 1985; Cumberledge & Carbon, 1987; Gaudet & Fitzgerald-Hayes, 1987). More recently, the presence of homopolymeric runs of A and T within the CDEII elements were identified to play a significant role in centromere function *in vivo* (Baker & Rogers, 2005), but the underlying mechanism requiring these homopolymeric runs remains unknown due to the inherent instability of these nucleosomes *in vitro* (Dechassa *et al*., 2011; Dechassa *et al*., 2014; Xiao *et al*., 2011). One possibility is that these sequences play a role in exclusion of the canonical H3 nucleosome, as H3 eviction has been proposed as a potential function of centromeres (Dechassa *et al*., 2011; Shukla *et al*, 2018). Due to their difficulty to study in cells, it remains unclear why these CDEII centromere sequences are essential *in vivo* yet are such poor templates for nucleosome formation and kinetochore assembly *in vitro*.

To resolve this paradox between the requirements for centromere sequence in nucleosome formation *in vitro* versus *in vivo,* it is imperative to determine what additional factors stabilize centromeric nucleosomes in a physiological context. Recent structural reconstitutions that contain a Cse4^CENP-A^ nucleosome in complex with additional inner kinetochore proteins (CCAN) have identified significant interactions between centromeric DNA and inner kinetochore proteins around the nucleosome (Guan *et al*., 2021; Yan *et al*, 2019). These reconstitutions have provided insight into potential candidate factors, yet the use of non-native centromeric DNA limits model testing. Unfortunately, studying the formation of centromeric nucleosomes in cells is extremely challenging because the centromeres are too close to be resolved individually by light microscopy and because they remain fully occupied for most of the cell cycle (Dhatchinamoorthy *et al*, 2017; Joglekar *et al*, 2006).

To address these limitations, we developed a new technique utilizing total internal <u>r</u>eflection fluorescent <u>m</u>icroscopy (TIRFM) to enable direct observation of centromeric nucleosome formation with single molecule resolution. Our approach was inspired by the “colocalization single molecule spectroscopy” (CoSMoS) studies of DNA transcription and RNA splicing complexes (Friedman *et al*, 2006; Friedman & Gelles, 2012; Hoskins *et al*, 2011). In our adaptation of the technique, individual centromere DNAs are linked sparsely to a glass surface and formation of single Cse4^CENP-A^ nucleosomes on the individual DNAs is observed in cell extract in real time. To achieve this resolution, *Saccharomyces cerevisiae* was used as a model system due to their simplified and sequence-defined point centromeres that contain only a single Cse4^CENP-A^ nucleosome (Cole *et al*., 2011; Furuyama & Biggins, 2007). Stable recruitment of Cse4^CENP-A^ was highly specific and dependent upon native centromere sequence, recapitulating *in vivo* requirements for nucleosome formation. Through continuous visualization of individual centromeres, we observed unexpectedly dynamic association of Cse4^CENP-A^. Specifically, we found that Cse4^CENP-A^ deposition occurred in two distinct steps: first, a “targeting” step, consisting of a reversible binding of Cse4^CENP-A^ for which Scm3^HJURP^ was dispensable, followed by a second “stabilization” step, for which Scm3^HJURP^ and DNA-binding inner kinetochore proteins were required. Stabilization was blocked by constraining both ends of the centromeric DNA template, suggesting that it also requires DNA wrapping around the Cse4^CENP-A^ nucleosome. Stabilization was also significantly influenced by both the sequence composition of the CDEII element and the subsequent binding of inner kinetochore proteins during kinetochore assembly. Together, these findings shed new light on the mechanisms that catalyze formation of a robust centromeric DNA-based platform for kinetochore assembly and provide a foundation to address additional steps in the kinetochore assembly process.

## Results

### Efficient recruitment of Cse4^CENP-A^ to individual centromeric DNAs

To study the requirements and dynamics of centromeric nucleosome assembly, we adapted our recently developed method for bulk assembly of yeast kinetochores *de novo* in cell extracts (Lang *et al*, 2018), modifying it for single molecule imaging via TIRFM. Template DNAs consisting of the chromosome III centromere (117 bp), with ∼70 bp of pericentromeric DNA plus ∼250 bp of linker DNA on each side (referred to as ‘CEN DNA’), were linked sparsely to a streptavidin-functionalized coverslip surface through biotin tags (Crawford *et al*, 2008) (Figure 1A). Dye-labels added to the free ends of the DNAs allowed their visualization at the single molecule level. Introducing whole cell extracts prepared from strains with fluorescent kinetochore proteins into the chamber enabled the recruitment and retention of the labeled kinetochore proteins on ∼1000 individual CEN DNA molecules to be monitored in a single field of view. Initially, we performed simple endpoint analyses, where we incubated cell lysate for 90 min with surface-linked CEN DNAs, washed the lysate from the chamber and then measured colocalization from individual images.

**Figure 1.**
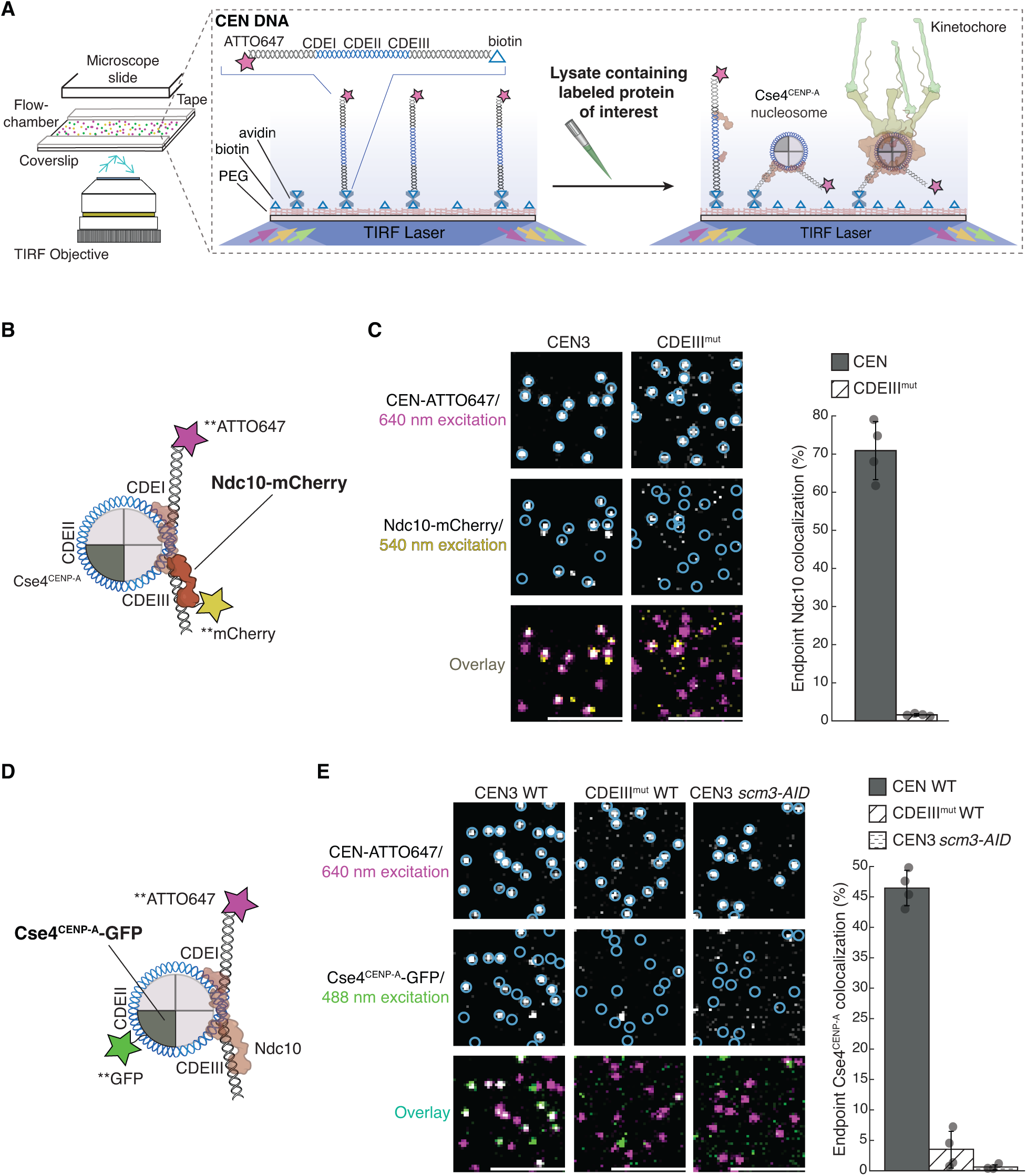
Ndc10 and Cse4^CENP-A^ assemble with high efficiency and specificity onto CEN DNAs in extract. A Schematic of the TIRFM colocalization assay. Yeast lysate containing a fluorescent protein(s) of interest is added to a coverslip with immobilized fluorescent CEN DNA. After incubation, the lysate is washed from the chamber and the CEN DNA and fluorescent kinetochore proteins are imaged via TIRFM. B Schematic of fluorescent label location around the centromeric nucleosome used in (C) for colocalization imaging. C Example images of TIRFM endpoint colocalization assays. Top panels show CEN DNA (Top-left panel, blue circles) or CDEIII^MUT^ CEN DNA (top-right panel, blue circles) visualized in lysates containing Ndc10-mCherry. Middle panels show the visualized Ndc10-mCherry on CEN DNA (middle-left panel) or CDEIII^MUT^ DNA (middle-right panel) with colocalization shown in relation to blue DNA circles. Bottom panels show overlay of DNA channel (magenta) with Ndc10-mCherry (yellow) on CEN DNA (bottom-left panel) or CDEIII^MUT^ DNA (bottom-right panel). Graph shows the quantification of Ndc10 endpoint colocalization on CEN DNA and on CDEIII^MUT^ CEN DNA (70 ± 7.6%, 1.6 ± 0.3% respectively, avg ± s.d. n=4 experiments, each examining ∼1,000 DNA molecules from different extracts). D Schematic of fluorescent label location around the centromeric nucleosome used in (E) for colocalization imaging. E Example images of TIRFM endpoint colocalization imaging. Top panels show CEN DNA (top-left panel, blue circles) or CDEIII^MUT^ CEN DNA (top-middle panel, blue circles) visualized in lysates that included Cse4^CENP-A^-GFP or CEN DNA in lysates that lacked Scm3^HJURP^ (*scm3-AID)* (top-right panel, blue circles). Middle panels show Cse4^CENP-A^ GFP visualized on CEN DNA (middle-left panel) or CDEIII^MUT^ CEN DNA (center panel) or on CEN DNA in lysates lacking Scm3^HJURP^ (*scm3-AID)* (middle-right panel) with colocalization shown in relation to blue DNA circles. Bottom panels show overlay of CEN DNA channel (magenta) with Cse4^CENP-A^-GFP (green) on CEN DNA (bottom-left panel) or CDEIII^MUT^ DNA (bottom-middle panel) or on CEN DNA in lysates lacking Scm3^HJURP^ (*scm3-AID)* (bottom-right panel). Scale bars 3μm. Graph shows quantification of observed colocalization of Cse4 on CEN DNA and on CDEIII^MUT^ CEN DNA or on CEN DNA in lysates that lacked Scm3^HJURP^ (47 ± 2.9%, 3.5 ± 3.0% and 0.6 ± 0.4% respectively, avg ± s.d. n=4 experiments, each examining ∼1,000 DNA molecules from different extracts).

We first tested whether Cse4^CENP-A^ and the CBF3 complex, which is required for Cse4^CENP-A^ localization *in vivo,* were specifically recruited to the individual CEN DNA molecules (Cho & Harrison, 2011a; Lang *et al*., 2018). Using extracts from cells expressing endogenously tagged Ndc10-mCherry, a CBF3 complex component (Figure 1B), we observed high endpoint colocalization of Ndc10 on CEN DNA (∼70%, Figure 1C left panel). To ensure specific colocalization, we tested a mutant CEN DNA template (CDEIII^mut^) with two substitutions that prevent CBF3 complex binding (Lang *et al*., 2018; McGrew *et al*, 1986). Ndc10-mCherry endpoint colocalization on this mutant template was nearly abolished (Figure 1C-right panel). We next sought to monitor endpoint colocalization of Cse4^CENP-A^-GFP using an internal Cse4 tag that is fully functional (Figure 1D), as it was previously shown that the position of the GFP tag in Cse4 affects protein function (Wisniewski *et al*, 2014). Cse4^CENP-A^-GFP showed high endpoint colocalization of ∼50% (Figure 1E-left panel). This localization was specific because it was nearly abolished on the CDEIII^mut^ DNA template (Figure 1E-middle panel), consistent with the CBF3 complex being required for Cse4^CENP-A^ centromere localization (Cho & Harrison, 2011a; Lang *et al*., 2018). Likewise, proteasomal degradation of the essential chaperone Scm3^HJURP^ also prevented Cse4^CENP-A^ from colocalizing to CEN DNAs (Lang *et al*., 2018; Nishimura *et al*, 2009) (Figure 1E-right panel). To further probe the fidelity of the assay, we quantified the stoichiometry of Ndc10 and Cse4^CENP-A^ via photobleaching assays. The Ndc10-mCherry that colocalized with CEN DNA photobleached predominantly in two steps (Appendix Figure S1A-B) with a step distribution similar to other previously characterized dimeric proteins (Popchock *et al*, 2018; Popchock *et al*, 2017). This is consistent with structural studies showing that Ndc10 is a homodimer within the CBF3 complex and with the range of stoichiometries reported *in vivo* (Cho & Harrison, 2011a; Guan *et al*., 2021; Joglekar *et al*., 2006; Leber *et al*, 2018; Wisniewski *et al*., 2014). The Cse4^CENP-A^ that colocalized with CEN DNA yielded similar results, with the majority of Cse4^CENP-A^-GFP photobleaching in two steps (Appendix Figure S1C-D), as expected based on nucleosome reconstitutions and consistent with fluorescence and photobleaching analyses of Cse4^CENP-A^ at centromeres *in vivo* (Aravamudhan *et al*, 2013; Camahort *et al*., 2007; Guan *et al*., 2021; Joglekar *et al*., 2006). Taken together, these results show that recruitment of Cse4^CENP-A^ onto individual CEN DNAs depends on the known requirements *in vivo* and suggest that the copy numbers of both Ndc10 and Cse4^CENP-A^ recruited onto the DNAs match their copy numbers at kinetochores *in vivo*.

### Cse4^CENP-A^ binds more stably when preceded by CBF3 complex component Ndc10

We next set out to monitor the dynamics of Ndc10 and Cse4^CENP-A^ centromere targeting by performing continuous time-lapse TIRFM. We tagged Ndc10 with mCherry in cells also expressing GFP-labeled Cse4^CENP-A^ to allow simultaneous detection in separate color channels (Figure 2A). To assist in the analysis of colocalization events taking place on hundreds of CEN DNA templates simultaneously, we developed an automated analysis software package in MATLAB (see Methods). Briefly, we collected a 45 min TIRFM time-lapse with acquisitions every 5 sec in the protein channels (488 nm and 568 nm) and every 60 sec in the DNA channel (647 nm). The software identified CEN DNAs and then subsequently monitored both protein channels to determine colocalization residence lifetimes at each CEN DNA (Figure 2A), enabling the rapid quantification of many independent colocalization events within a single field of view. We applied this analysis to simultaneously measure residence lifetimes for both Ndc10 and Cse4^CENP-A^ (Figure 2A, Figure EV1A). Kaplan-Meier analysis of the estimated survival functions showed that Cse4^CENP-A^ had a shorter median lifetime (approximately half) than the median Ndc10 lifetime (Figure 2B). We note that occasionally one of the proteins was present when imaging began or persisted until the acquisition ended (censored events). To avoid the potential exclusion of long-lived residences and because these censored events were typically <5% of the total residences observed, they were included in this analysis. To ensure that the dynamic Cse4^CENP-A^ behavior we observed was not a consequence of a particular CEN sequence, we performed residence lifetime assays on several CEN templates from different chromosomes. The median Cse4^CENP-A^ lifetimes were similar between varying chromosome CEN templates (103 s, 103 s, and 109 s), all shorter than the median survival time of Ndc10 (Appendix Figure S2A). Consistent with this, we also confirmed similar Cse4^CENP-A^ endpoint colocalization recruitment on several different CEN templates (Appendix Figure S2B, C). During this analysis, we found that the Cse4^CENP-A^-CEN DNA endpoint colocalization was lower in genetic backgrounds where Cse4^CENP-A^ and another kinetochore protein were fluorescently tagged compared to when only Cse4^CENP-A^ was tagged, likely due to mild genetic interactions between tagged proteins. However, this did not prevent analyses of the arrival times and residence lifetimes of the tagged components relative to one another.

**Figure 2.**
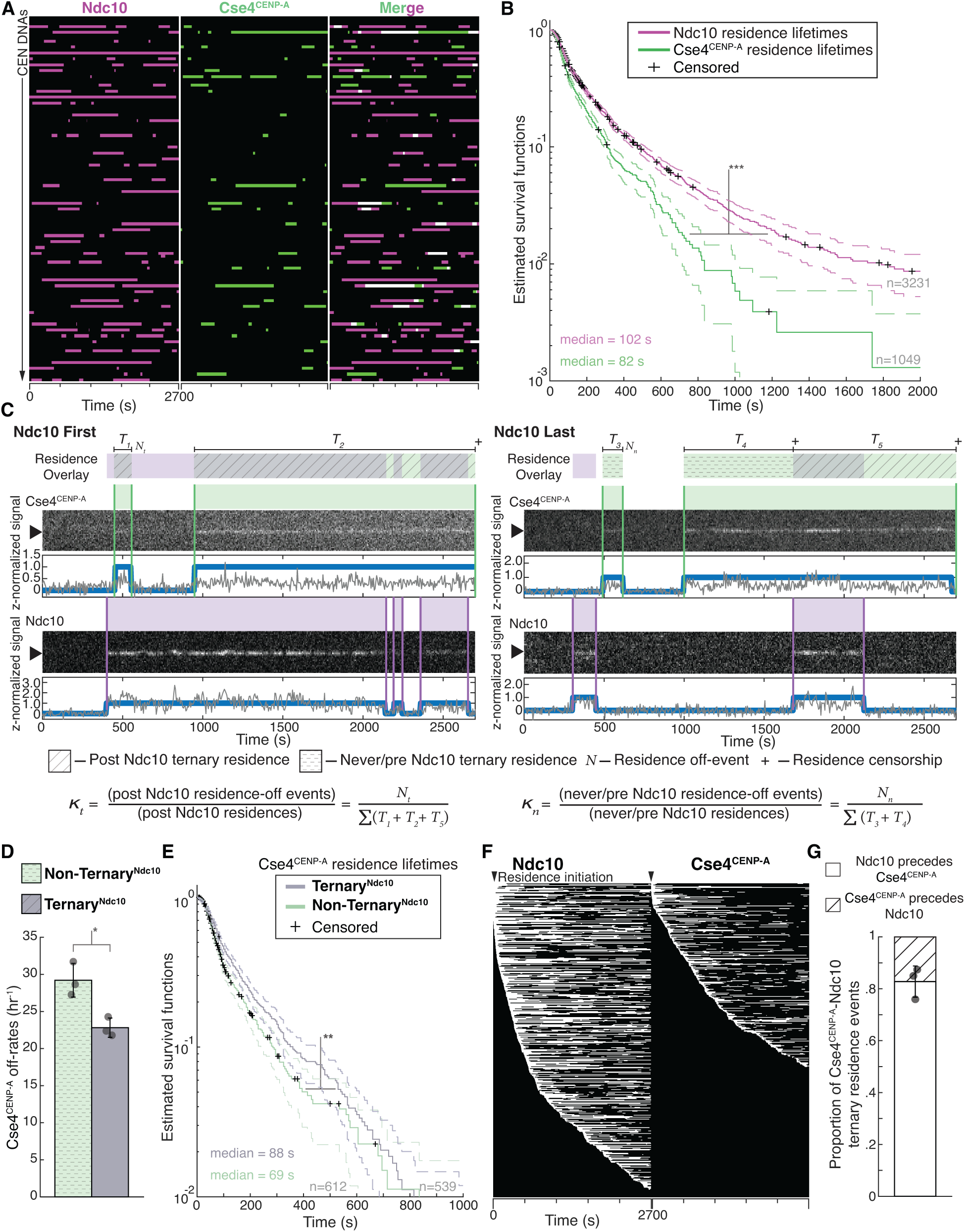
Ndc10 stabilizes Cse4^CENP-A^ on CEN DNA. A Example graph of total identified colocalization residences observed on CEN DNA per imaging sequence. Each row represents one identified CEN DNA with all identified residences shown over entire imaging sequence (2700 s) for Ndc10 (magenta) and Cse4^CENP-A^ (green). Instances where Ndc10 and Cse4^CENP-A^ residence coincides (white-merge) represent simultaneous observation of both proteins on single CEN sDNA, termed ternary residences. Complete series shown in Figure EV1A. B Estimated survival function plots of Kaplan-Meier analysis of CEN DNA residence lifetimes of Ndc10 (magenta – median lifetime of 102 s, n=3231 over 3 experiments of ∼1000 DNA molecules using different extracts) and Cse4^CENP-A^ (red – median lifetime of 79 s, n=1049 over 3 experiments of ∼1000 DNA molecules using different extracts). Significant difference between ternary Ndc10 residence lifetime survival plots (***) compared to Cse4^CENP-A^ (two-tailed p-value of 0 as determined by log-rank test). 95% confidence intervals indicated (dashed lines), right-censored lifetimes (plus icons) were included and unweighted in survival function estimates. C Schematic for off-rate estimate calculations with representative residence lifetime assay traces of Ndc10 and Cse4^CENP-A^ on CEN DNA. Example kymographs of Cse4^CENP-A^ (top-488 nm) and Ndc10 (bottom-568 nm) in relation to a single identified CEN DNA (arrow), with normalized intensity trace below (grey) as well as identified colocalization residences (blue). <u>Ndc10 First</u> example (left) illustrates Ndc10 colocalization that precedes Cse4^CENP-A^, yielding only **Ternary^Ndc10^**Cse4^CENP-A^ residences. <u>Ndc10 Last</u> example (right) illustrates Cse4^CENP-A^ colocalization that precedes Ndc10, yielding both **Non-Ternary^Ndc10^** Cse4^CENP-A^ residences and **Ternary^Ndc10^** Cse4^CENP-A^ residence after Ndc10 colocalization initiation. Images were acquired every 5 seconds with normalized fluorescence intensity shown in arbitrary units. Bottom panel includes formulas used to calculate the ternary Cse4^CENP-A^ off-rate, *k_t_*, where *N_t_* represents the total number of apparent Cse4-GFP detachments recorded during ternary residences, and ∑(*T_1_+T_2_+T_5_*) represents the total amount of ternary residence time observed. As well as the formula for non-ternary Cse4 off-rate, *k_n_*, where *N_n_* represents the total number of apparent Cse4 GFP detachments recorded during non-ternary residences, and ∑(*T_3_+T_4_*) represents the total amount of non-ternary residence time observed. D Cse4^CENP-A^ has slower off-rates after colocalization with Ndc10 on CEN DNA. Quantification of the estimated Cse4^CENP-A^ off-rates of the **Non-Ternary^Ndc10^** and **Ternary^Ndc10^** pools (29 hr^-1^ ± 2 hr^-1^ and 22 hr^-1^ ± 1 hr^-1^ respectively, avg ± s.d. n=1151 over 3 experiments of ∼1000 DNA molecules using different extracts). Significant difference between off-rates (*) with a P-value of .02 as determined by two-tailed unpaired *t*-test. E Kaplan-Meier analysis of **Ternary^Ndc10^** residence lifetimes of Cse4^CENP-A^ on CEN DNA (purple – median lifetime of 88 s, n=539 over 3 experiments of ∼1000 DNA molecules using different extracts) and of **Non-Ternary^Ndc10^** Cse4^CENP-A^ residence lifetimes (teal – median lifetime of 69 s, n=612 over 3 experiments of ∼1000 DNA molecules using different extracts). There is a significant difference (**) between **Ternary^Ndc10^** and **Non Ternary^Ndc10^** lifetime survival plots (two-tailed p-value of 7.4e-4 as determined by log rank test). 95% confidence intervals indicated (dashed lines), right-censored lifetimes (plus icons) were included and unweighted in survival function estimates. F Example plot of total identified residences observed on CEN DNA per imaging sequence of Ndc10 (left) and Cse4^CENP-A^ (right) independently sorted by residence initiation time, starting at the top with initiation time of 0 s (arrow). G Timing of all non-simultaneous Ndc10 and Cse4^CENP-A^ ternary residence events. The proportion when Ndc10 precedes Cse4^CENP-A^ is 0.83 ± .06 and when Cse4^CENP-A^ precedes Ndc10 is 0.17 ± .06 (avg ± s.d., n=504 over 3 experiments of ∼1000 DNA molecules using different extracts). Simultaneous arrival defined as residence initiation within 5 s of each other and comprise .07 ± .04 of all ternary residence events (avg ± s.d., n=35 over 3 experiments of ∼1000 DNA molecules using different extracts).

The dynamic behavior of Cse4^CENP-A^ was surprising since it has been reported to be stably bound to centromeres *in vivo* outside of S phase (Joglekar *et al*, 2008; Wisniewski *et al*., 2014). However, the transient Cse4^CENP-A^-CEN DNA interactions that we readily detected in our single molecule assays would be undetectable *in vivo* due to reduced signal-to-noise and poorer time resolution. We considered two factors that could contribute to the dynamic Cse4^CENP-A^ behavior we observed. First, the cellular extracts could contain negative regulatory factors, such as chromatin remodelers that counter nucleosome formation. To test this, we removed the lysate and analyzed protein stability on the CEN DNA. Remarkably, Cse4^CENP-A^ and Ndc10 exhibited ∼70% retention on CEN DNA after 24 hrs incubation in buffer at 25 °C (Figure EV1B, C), suggesting there are factors in the lysate that actively disrupt the centromeric nucleosome. Second, we considered the possibility that photobleaching could make Cse4^CENP-A^ appear dynamic. To address this, we estimated the photo-stability of the fluorophores by measuring their respective lifetimes under our imaging parameters and found that the Ndc10 residence lifetimes (Figure EV1D) were more potentially limited by photo-stability than the Cse4^CENP-A^ lifetimes (Figure EV1E), the majority of which were not truncated by photobleaching. Despite photobleaching limiting some residence lifetimes, the relationships we observed in residence lifetime assays were consistent with trends observed in the endpoint colocalization assays. For example, the longer median lifetime of Ndc10 compared to Cse4^CENP-A^ (102 s vs. 79 s) correlated with higher endpoint colocalization (70% vs. 47%). Taken together, these data suggest that Cse4^CENP-A^ is initially more dynamic than Ndc10 at centromeres, prior to its stable incorporation.

To further dissect the lifetime differences between Ndc10 and Cse4^CENP-A^, we sought to determine if the presence of Ndc10 affected the behavior of Cse4^CENP-A^ on CEN DNA. To do this, we identified instances where a Cse4^CENP-A^ residence coincided with Ndc10 at any time on the same CEN DNA, termed “ternary residence”. To assess whether the formation of a ternary^Ndc10^ residence altered Cse4^CENP-A^ stability on CEN DNA, we sought to estimate the off-rates of Cse4^CENP-A^ in the ternary versus non-ternary contexts. Although 84% of the total Cse4^CENP-A^ ternary residence time had Ndc10 associated, we wanted to rigorously ensure there was no selection bias for long-lived Cse4^CENP-A^ residences because they are more likely to form ternary associations. We therefore further divided the ternary^Ndc10^ Cse4^CENP-A^ residences according to the timing of Ndc10 association. When Ndc10 arrived prior to or simultaneously with Cse4^CENP-A^, we included the entire residence time (Ndc10 first - T_1_, T_2_, Figure 2C). In the cases where Cse4^CENP-A^ association preceded Ndc10, we only included the residence time after Ndc10 arrived (Ndc10 last - T_5_, Figure 2C). Because we could not differentiate between Ndc10 turnover or photobleaching, we included the entirety of a continuous Cse4^CENP-A^ residence pulse after initial ternary association with Ndc10, regardless of Ndc10 residence length or subsequent associations. For the non-ternary^Ndc10^ residences (T_3_, T_4_, Figure 2C), we included any Cse4 residence prior to the arrival of Ndc10 as well as Cse4^CENP-A^ residences where Ndc10 was never present.

To estimate off-rates for ternary and non-ternary Cse4^CENP-A^, we counted the numbers of off-events observed in each category, N_t_ and N_n_ respectively, and divided each by the total observed times from each residence pool, T_t_ = SUM(of all T_1_, T_2_, and T_5_) to calculate the off-rate *k_t_*(Figure 2C) and T_n_ = SUM(of all T_3_ and T_4_) to calculate the off-rate *k_n_* (Figure 2C). The off-rate for ternary^Ndc10^ Cse4^CENP-A^ was significantly lower than for non-ternary^Ndc10^, 22/hr vs 29/hr, indicative of stabilization of Cse4^CENP-A^ on CEN DNA after ternary Ndc10 association (Figure 2D). We also performed Kaplan-Meier analysis to estimate their survival lifetimes and found a significant increase in the median lifetime of ternary^Ndc10^ compared to non-ternary^Ndc10^ residences (88 s vs. 69 s, Figure 2E). Consistent with this stabilization, the proportion of Cse4^CENP-A^ residences that were long lived (>300s) after ternary^Ndc10^ association increased approximately two-fold (Figure EV1F). While we did not estimate on-rates, comparison of Ndc10 and Cse4^CENP-A^ residence pulses when organized by their initiation times showed more rapid association of Ndc10 to CEN DNA (Figure 2F). This potential for differing on-rates may explain why when ternary associations were formed, Ndc10 typically preceded Cse4^CENP-A^ arrival to the CEN DNA (Figure 2G). We note that it was surprising to see Cse4^CENP-A^ at the CEN DNA in the absence of Ndc10. To determine whether Ndc10 is present but undetectable in these instances, we analyzed Cse4^CENP-A^ lifetimes on the mutant CDEIII^mut^ CEN DNA template which abolishes Ndc10 association. Although overall Cse4^CENP-A^ associations were reduced, we observed transient Cse4^CENP-A^ associations with significantly reduced residence lifetimes on CDEIII^mut^ CEN DNA, suggesting that Cse4 has some intrinsic CEN DNA binding in the absence of CBF3 (Figure EV1G, H). Taken together, these observations are consistent with *in vivo* findings, where it has been established that Ndc10 is required for Scm3^HJURP^-dependent Cse4^CENP-A^ deposition, and where Ndc10 is thought to coordinate with the Cbf1 complex to promote nucleosome formation (Cho & Harrison, 2011a).

### The chaperone Scm3^HJURP^ is a limiting cofactor that promotes stable Cse4^CENP-A^ association

The biphasic Cse4^CENP-A^ localization behavior, with many short colocalizations (<120 s) and fewer long colocalizations (>300 s), was not fully correlated with Ndc10 occupancy, as 60% of ternary^Ndc10^ Cse4^CENP-A^ residences were still short-lived. We therefore further investigated this behavior by simultaneously visualizing Cse4^CENP-A^ and its essential chaperone protein Scm3^HJURP^, which exhibits DNA-binding activity and is required for Cse4^CENP-A^ centromere localization in cells (Camahort *et al*., 2007; Mizuguchi *et al*., 2007; Stoler *et al*., 2007; Xiao *et al*., 2011) (Figure 3A). Residence lifetimes for Scm3^HJURP^ on CEN DNA were shorter than Cse4^CENP-A^ lifetimes (Figure 3A, B, Figure EV2A, B), consistent with its faster turnover rate in vivo (Wisniewski et al., 2014). To estimate the fraction of time Scm3^HJURP^ and Cse4^CENP-A^ colocalized together on CEN DNA, we analyzed the ternary^Scm3^ residence times. We found that Scm3^HJURP^ co occupied the CEN DNA an average of 56.0% (n=393) of the total Cse4^CENP-A^ residence time, a lower occupancy rate than that of Ndc10 and Cse4^CENP-A^ (84.3%), as expected given its more rapid turnover. Strikingly, we discovered a high proportion of total Cse4^CENP-A^ residences that occurred without Scm3^HJURP^ (0.78 vs. 0.22, Figure 3C), which was unexpected because Scm3^HJURP^ is thought to be required to target Cse4^CENP-A^ to centromeres *in vivo* (Camahort *et al*., 2007; Stoler *et al*., 2007).

**Figure 3.**
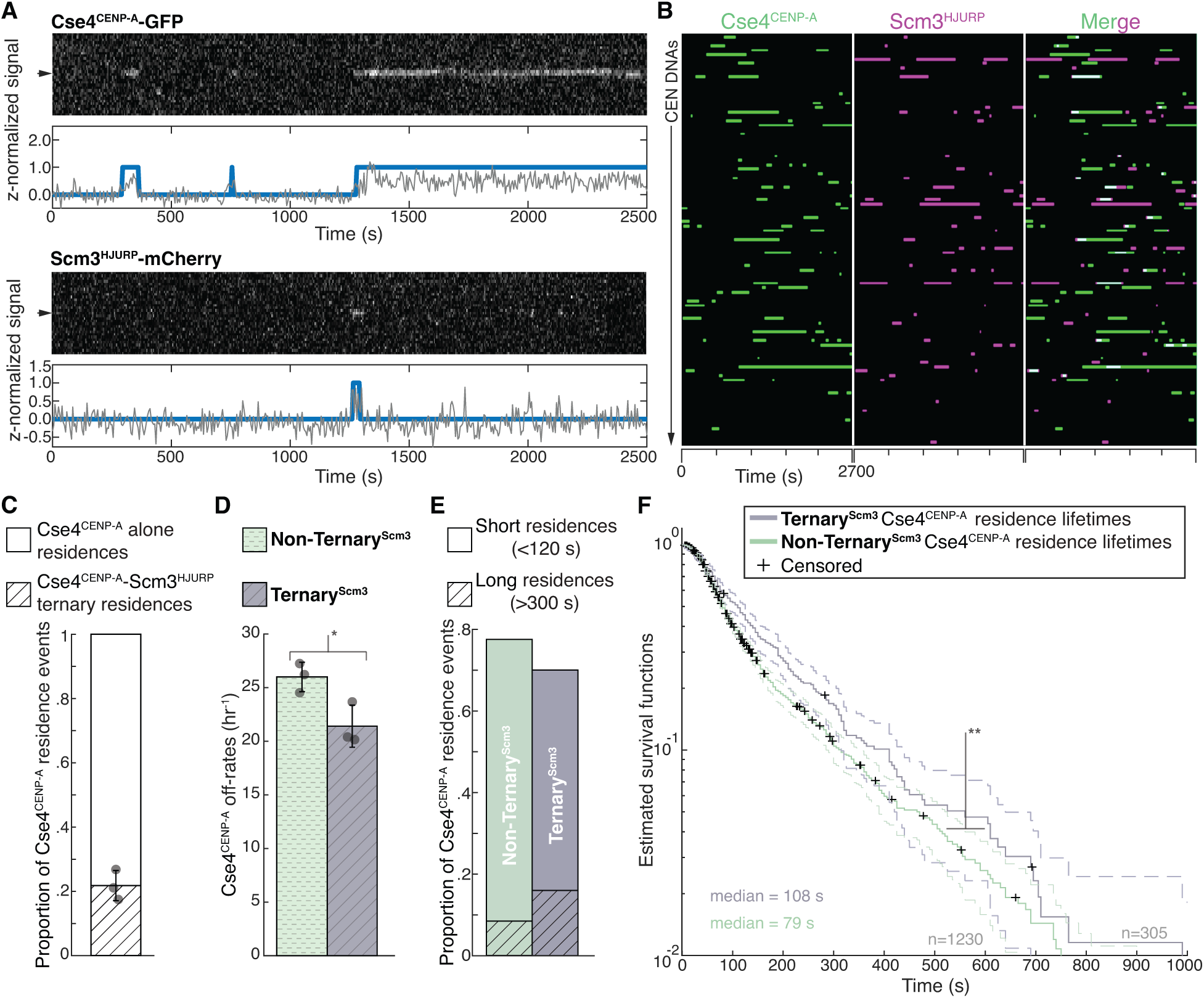
Cse4^CENP-A^ residence time on centromeric DNA is increased in the presence of its chaperone Scm3^HJURP^. A Representative residence lifetime assay traces of Cse4^CENP-A^ and Scm3^HJURP^ on a single CEN DNA. Top panel includes kymograph of Cse4^CENP-A^ (top-488 nm) in relation to single identified CEN DNA (arrow), with normalized intensity trace (grey-bottom) as well as identified residences (blue). Bottom panel includes kymograph of Cse4^CENP-A^ (bottom-568 nm) in relation to the same identified CEN DNA (arrow), with normalized intensity trace (grey-bottom) as well as identified residences (blue). Images acquired every 5 seconds with normalized fluorescence intensity shown in arbitrary units. B Example plot of total identified residences observed on CEN DNA per imaging sequence. Each row represents one identified CEN DNA with all identified residences shown over entire imaging sequence (2700 s) for Cse4^CENP-A^ (green) and Scm3^HJURP^ (magenta). Cases where Scm3^HJURP^ and Cse4^CENP-A^ residence coincides represent simultaneous observation of both proteins on single CEN DNA, termed ternary residence (merge, white). Complete plot shown in Figure EV2A. C Quantification of the proportion of Cse4^CENP-A^ and Scm3^HJURP^ ternary residences with CEN DNA compared to CEN DNA residences of Cse4^CENP-A^ alone (0.22 ± .05 and 0.78 ± .05 respectively, avg ± s.d. n=1419 over 3 experiments of ∼1000 DNA molecules using different extracts). Ternary residences include simultaneous Scm3^HJURP^ residence at any point during continuous Cse4^CENP-A^ residence on CEN DNA. D Quantification of the estimated off-rates of Cse4^CENP-A^ that never formed a ternary residence (**Non-Ternary^Scm3^**) and of Cse4^CENP-A^ after ternary residence with Scm3^HJURP^ (**Ternary^Scm3^**) on CEN DNA (26 hr^-1^ ± 1 hr^-1^ and 21 hr^-1^ ± 2 hr^-1^ respectively, avg ± s.d. n=1535 over 3 experiments of ∼1000 DNA molecules using different extracts). Significant difference between off-rates (*) with a P-value of .03 as determined by two-tailed unpaired *t*-test. E Quantification of the proportion of short residences (<120 s) and long residences (>300 s) of Cse4^CENP-A^ alone or in ternary residences on CEN DNA. Short and long **Non Ternary^Scm3^** Cse4^CENP-A^ residences (.69 and .08 respectively, n=1063 over 3 experiments of ∼1000 DNA molecules using different extracts) and short and long **Ternary^Scm3^** Cse4^CENP-A^ residences (.54 and .16 respectively, n=305 over 3 experiments of ∼1000 DNA molecules using different extracts). F Longer residence lifetimes are measured for Cse4^CENP-A^ after Scm3^HJURP^ colocalizes. Estimated survival function plots of Kaplan-Meier analysis of the lifetimes of **Ternary^Scm3^**Cse4^CENP-A^ residences on CEN DNA (purple – median lifetime of 108 s, n=305 over 3 experiments of ∼1000 DNA molecules using different extracts) and **Non-Ternary^Scm3^** Cse4^CENP-A^ residences on CEN DNA (green of 79 s, n=1230 over 3 experiments of ∼1000 DNA molecules using different extracts). Significant difference (**) between **Ternary^Scm3^** and **Non-Ternary^Scm3^ lifetime survival plots** (two-tailed p-value of 2.1e-3 as determined by log-rank test). 95% confidence intervals indicated (dashed lines), right censored lifetimes (plus icons) were included and unweighted in survival function estimates.

We next assayed whether the two distinct Cse4^CENP-A^ residence subpopulations, ternary^Scm3^ and non-ternary^Scm3^, behaved differently by quantifying their estimated off rates as described for Ndc10 (Figure 2C). We found a significant decrease in the average off-rate of ternary^Scm3^ Cse4^CENP-A^ residences when compared to non-ternary^Scm3^ (26 hr^-1^ vs. 21 hr^-1^, Figure 3D). Consistent with this observed stabilization of Cse4^CENP-A^ after Scm3^HJURP^ association and the requirement for Scm3^HJURP^ to localize Cse4^CENP-A^ to centromeres in vivo, the proportion of Cse4^CENP-A^ residences that were long lived (>300s) increased approximately two-fold among the ternary^Scm3^ events (Figure 3E). We also performed Kaplan-Meier analysis to estimate their survival lifetimes and found a significant increase in the median lifetime of ternary^Scm3^ residences when compared to non-ternary^Scm3^ (108 s vs. 79 s, Figure 3F). Together, these results are consistent with Scm3^HJURP^ stabilizing Cse4^CENP-A^ at centromeres and then rapidly exchanging after Cse4^CENP-A^ incorporation (Wisniewski *et al*., 2014).

To further dissect the role of Scm3^HJURP^ in Cse4^CENP-A^ centromere localization, we calculated the off-rates for Scm3^HJURP^ in ternary^Cse4^ and non-ternary^Cse4^ contexts. Unlike Cse4^CENP-A^, Scm3^HJURP^ was not significantly stabilized after ternary association with Cse4^CENP-A^, as the off-rates and median lifetimes of both contexts were similar (Figure EV2C, D). These findings are consistent with a model of Scm3^HJURP^ turnover after association with Cse4^CENP-A^. Next, we assayed the relative arrival order of Cse4^CENP-A^ versus Scm3^HJURP^ on CEN DNA. Contrary to models that suggest the chaperone delivers Cse4^CENP-A^ to centromeres, only ∼20% of the ternary^Scm3^ residences resulted from co arrival of Cse4^CENP-A^ and Scm3^HJURP^ (Figure EV2E). The remaining ternary associations consisted of 46% where Scm3^HJURP^ preceded Cse4^CENP-A^ and 34% where Cse4^CENP-A^ preceded Scm3^HJURP^ (Figure EV2E). Residence lifetimes of these differing ternary pools were similar, with no significant difference in estimated survival plots (Figure EV2F). This is consistent with our findings that Cse4^CENP-A^ can interact with the centromere in the absence of Scm3^HJURP^ (Figure 3C) and suggests that Scm3^HJURP^ catalyzes stable Cse4^CENP-A^ incorporation at centromeric DNA instead of delivering it to the centromere. Cse4^CENP-A^ centromere targeting is thought to occur through Scm3^HJURP^ binding to Ndc10, so we next monitored their behavior on mutant CDEIII^mut^ CEN DNA that lacks Ndc10 in lifetime residence assays (Figure EV3A). Strikingly, Scm3^HJURP^ residences were significantly increased when compared to those on CEN DNA (Figure EV3B), suggesting its intrinsic DNA-binding activity may be the primary driver of its CEN DNA binding and that stable Cse4^CENP-A^ association is required for its turnover (Xiao *et al*., 2011). In contrast, Cse4^CENP-A^ residences on CDEIII^mut^ CEN DNA had a much shorter median survival time compared to its CEN counterpart (Figure EV2C), consistent with Ndc10 and Scm3^HJURP^ stabilizing its association. In addition, the stabilization of Cse4^CENP-A^ after Scm3^HJURP^ association that was observed on CEN DNA (Figure 3D) was lost, as the lifetimes of ternary^Scm3^ and non-ternary^Scm3^ residences on CDEIII^mut^ CEN were similar (Figure EV3C). Taken together, these data are consistent with a requirement for Ndc10 in the localization of Cse4^CENP-A^ but not necessarily through Scm3 recruitment. One possibility is there is cooperativity between Ndc10 and Scm3^HJURP^ and both DNA-binding proteins must be present simultaneously at CEN DNA to promote stable Cse4^CENP-A^ incorporation.

Our finding that there was a population of Cse4^CENP-A^ lacking its chaperone led us to test whether the chaperone is limiting for stable Cse4^CENP-A^ centromere localization. To do this, we modified the availability of the Cse4^CENP-A^/Scm3^HJURP^ complex *in vivo* to assess resulting changes in Cse4^CENP-A^ behavior. The E3 ubiquitin ligase Psh1 and the chaperone Scm3^HJURP^ directly compete for Cse4^CENP-A^ binding (Zhou *et al*, 2021). We therefore reasoned that the amount of Cse4^CENP-A^-Scm3^HJURP^ complex in cells should be reduced by overexpressing Psh1 and, conversely, that the amount of Cse4^CENP-A^-Scm3^HJURP^ complex might be increased by overexpressing Scm3^HJURP^. To test these predictions, we introduced an ectopic copy of either Psh1 or Scm3^HJURP^ under an inducible *GAL* promoter into cells containing labeled Cse4^CENP-A^ and Scm3^HJURP^. Inducing short pulses of either protein did not significantly alter the total levels of Cse4^CENP-A^ in whole cell extracts (Figure EV4A). However, a short pulse of Scm3^HJURP^ overexpression did lead to a significant increase in the endpoint colocalization of Cse4^CENP-A^ on CEN DNA, while overexpression of Psh1 had the opposite effect (Figure EV4B, C). The median lifetime of Cse4^CENP-A^ on CEN DNA was also significantly increased when Scm3^HJURP^ was overexpressed and significantly decreased when Psh1 was overexpressed (Figure EV4D). These lifetime changes correlated with the propensity for Cse4^CENP-A^ to form ternary residences with Scm3^HJURP^ on the CEN DNA (Figure EV4E), suggesting they reflect changes in the availability of the complex. To assess if these perturbations to Cse4^CENP-A^ were consequential in cells, we grew cells under constant induction and found significant growth phenotypes when the Cse4^CENP-A^-Scm3^HJURP^ complex was limited (Figure EV4F). These data indicate that Scm3^HJURP^ is limiting for stable centromeric nucleosome formation in our single molecule assembly assay and suggest that availability of the Cse4^CENP-A^-Scm3^HJURP^ complex is likewise important for cell viability.

### Stable Cse4^CENP-A^ association is blocked when centromere DNAs are tethered at both ends

Our observation of two subpopulations of Cse4^CENP-A^ molecules on CEN DNAs, including brief binders and longer-residing molecules that correlated with Scm3^HJURP^, suggested a two-stage process for Cse4^CENP-A^ deposition that begins with transient, chaperone-independent targeting and is followed by chaperone-dependent stabilization. We hypothesized that the transition to more stable Cse4^CENP-A^ binding might require wrapping of the CEN DNA around the Cse4^CENP-A^-containing histone octamer. If wrapping were indeed required for stabilization, then restricting the ability of the DNA to adopt a wrapped configuration should inhibit its stabilization on CEN DNA. To test this prediction, we created CEN DNA templates with biotins located at both ends to enable double-ended attachment to the coverslip surface. Tethering both ends of the template should limit its ability to twist or shorten, and thus to encircle a Cse4^CENP-A^ histone octamer, behaviors which are proposed to be essential for Cse4^CENP-A^ nucleosome formation on centromeric DNA (Furuyama & Henikoff, 2009; Guo *et al*, 2021). To allow both ends to be tethered simultaneously to the coverslip surface, an additional biotin handle was introduced (instead of an organic dye) onto a shortened 250 bp version of the CEN DNA (Figure 4A). We first tested for a nucleosome-protected DNA fragment using a bulk assembly assay, in which template DNAs are linked to streptavidin-coated magnetic beads, incubated with cell extract to allow kinetochore assembly, and then washed, as done previously (Lang *et al*., 2018). In addition to the double-tethered template, controls using a single-tethered CEN template of the same length as well as a mutant CDEIII^mut^ template were also performed. After the assembly reactions, a mild micrococcal nuclease (MNase) digestion was performed to preferentially cut the unbound DNA in-between any assembled nucleosomes. Subsequent analysis of the DNA by gel electrophoresis showed strong protection of ∼140 bp of the single-tether template, but almost no protection of the double tethered template (Figure 4B). The CDEIII^mut^ template, which does not stably recruit Cse4^CENP-A^ (Lang *et al*., 2018), was completely unprotected as expected. Protection of the single-tether template was presumably due to formation of Cse4^CENP-A^-containing nucleosomes, because this DNA fails to recruit H3 (Lang *et al*., 2018). These observations suggest that spatial restriction of the template can prevent Cse4 nucleosome assembly.

**Figure 4.**
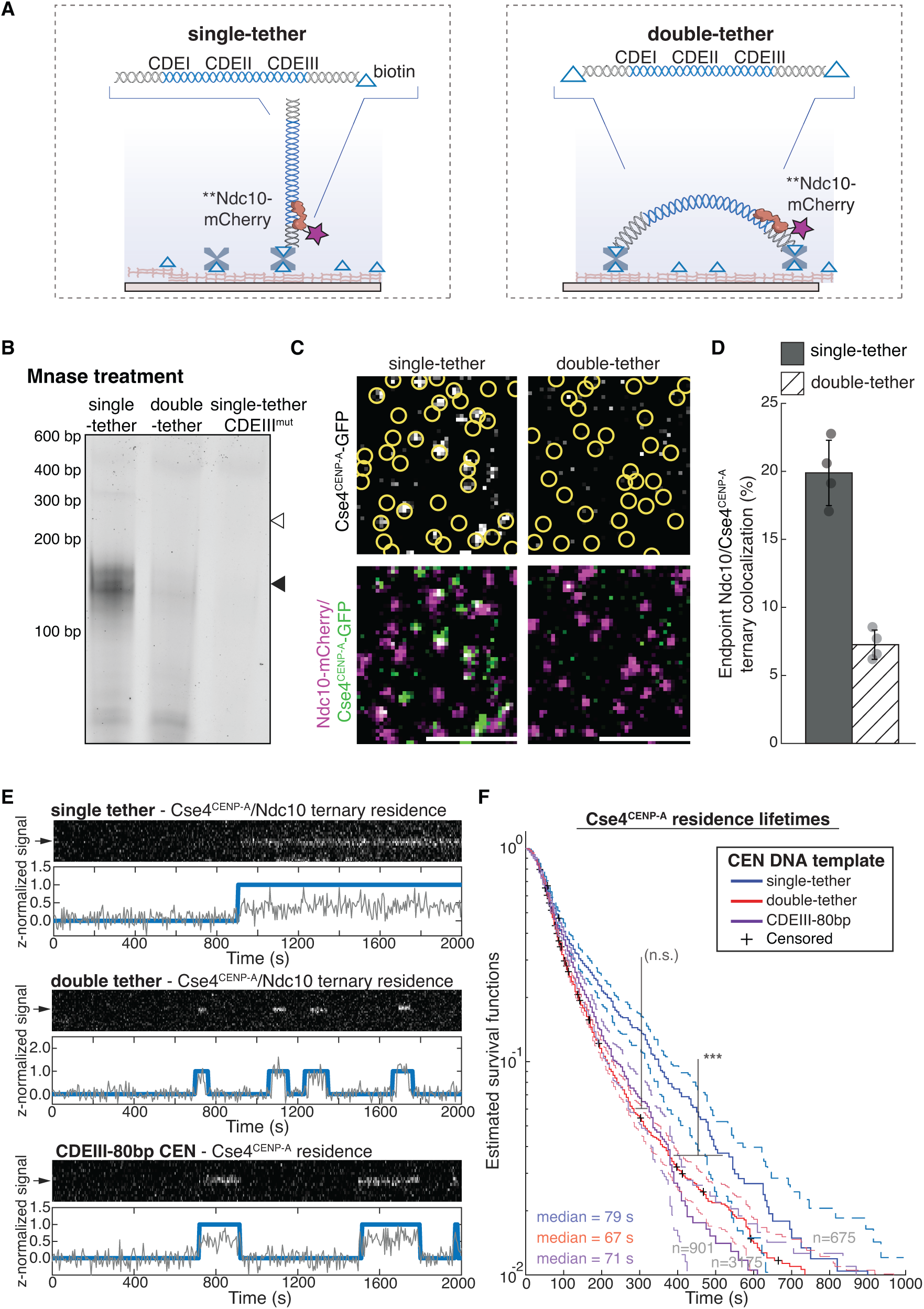
Synthetic restriction of nucleosome formation severely restricts Cse4^CENP-A^ residence lifetimes on CEN DNA. A Schematic of single vs. double-tether CEN DNA TIRFM colocalization assay. B Kinetochores were assembled on beads containing either a 250 bp single-tether, a 250 bp double-tether or a 250 bp CDEIII^mut^ CEN DNA. They were then treated with MNase, and the remaining DNA was visualized on an agarose gel. Black arrow indicates Cse4^CENP-A^ nucleosome protected DNA (∼150 bp); white arrow indicates theoretical location of undigested template DNA (250 bp). C Example images of TIRFM endpoint colocalization assays. Top panels show Cse4^CENP-A^/Ndc10 ternary colocalizations visualized on single-tethered CEN DNA (top left panel) or on double-tethered CEN DNA (top-right panel) with colocalization shown in relation to Ndc10 in yellow circles. Bottom panels show overlay of Ndc10 channel (magenta) with Cse4^CENP-A^ (green). Scale bars 3 μm. D Quantification of observed ternary colocalization of Cse4^CENP-A^ with Ndc10 (right) on single-tethered CEN DNA containing Ndc10 (19.9± 2.4%, avg ± s.d. n=4 experiments, each examining ∼1,000 DNA molecules from different extracts) and on double-tethered CEN DNA (7.3 ± 1.1%, avg ± s.d. n=4 experiments, each examining ∼1,000 DNA molecules from different extracts). E Representative residence lifetime assay traces of ternary Cse4^CENP-A^ residences with Ndc10 on single-tethered CEN DNA (top), double-tethered CEN DNA (middle) or of Cse4^CENP-A^ residence on CDEIII-80 bp CEN DNA (bottom). Each example includes kymographs of Cse4^CENP-A^ (488 nm-top) with normalized intensity trace (grey-bottom) as well as identified residences (blue). Images acquired every 5 seconds with normalized fluorescence intensity shown in arbitrary units. F Residence times for Cse4^CENP-A^ on double-tethered CEN DNAs are shorter than on single-tethered CEN DNAs and equivalent to those on non-functional mutant CEN DNAs. Estimated survival function plots of Kaplan-Meier analysis of ternary residences of Cse4^CENP-A^ with Ndc10 on single-tethered CEN DNA (blue – median lifetime of 79 s, n=675 over 3 experiments of ∼1000 DNA molecules using different extracts), on double tethered CEN DNA (red – median lifetime of 67 sec, n=3175 over 3 experiments of ∼1000 DNA molecules using different extracts) or Cse4^CENP-A^ lifetimes on CDEIII-80 bp CEN DNA (purple – median lifetime of 71 sec, n=901 over 3 experiments of ∼1000 DNA molecules using different extracts). No significant difference (n.s.) between double tethered 250 bp CEN DNA and 80 bp CDEIII CEN DNA lifetime survival plots (two-tailed p-value of 0.06 as determined by log-rank test). Significant difference (***) between double-tethered 250 bp CEN DNA and 250 bp single-tether DNA lifetime survival plots (two-tailed p-value of 1.4e-10 as determined by log-rank test). 95% confidence intervals indicated (dashed lines), right-censored lifetimes (plus icons) were included and unweighted in survival function estimates.

We next sought to analyze Cse4^CENP-A^ behavior on individual double-tethered CEN DNA templates in our TIRFM assay. Because the double tethered templates lacked an organic dye for visualization, we used Ndc10-mCherry as a fiducial marker for the CEN DNAs due to its high colocalization percentage (Figure 1C), rapid arrival, stable colocalization behavior, and our finding that most Cse4^CENP-A^ colocalizations occur after Ndc10 (Figure 2F, G). Endpoint colocalization of Cse4^CENP-A^ was reduced more than 2 fold on the double-tethered CEN DNAs relative to single-tethered controls (Figure 4C, D). Likewise, time-lapse experiments revealed that median Cse4^CENP-A^ residence lifetimes on double-tethered CEN DNAs were significantly shorter than on single-tethered templates (79 s vs. 67 s, Figure 4E, F). Median lifetimes on single-tethered templates were comparable to those of the previously measured ternary^Ndc10^ Cse4^CENP-A^ residences (79 s vs. 88 s, Figure 4E, F, Table 1). But median lifetimes of Cse4^CENP-A^ on double-tethered templates were significantly shorter, and closer to non-ternary^Ndc10^ Cse4^CENP-A^ residences (67 s vs. 69 s, Figure 4E, F, Table 1). As an alternative to physical restriction to prevent CEN DNA wrapping of Cse4^CENP-A^, we designed a dye-labeled and single-tethered, severely truncated 80-bp template (CDEIII-80bp CEN), that is too short to form a single wrap around the Cse4^CENP-A^-containing core particle (Figure EV5A, B). Cse4^CENP-A^ residences were transient on CDEIII-80bp CEN DNA (Figure EV5C) with a shortened median lifetime that was comparable to the median lifetime of ternary residences on double-tethered templates (71 s vs. 67 s, Figure 4E, F, Figure EV5D). The median lifetimes of Cse4^CENP-A^ residences on either mutant CEN DNA (CDEIII-80bp or double tethered) were closer to the median lifetime of non-ternary^Scm3^ Cse4^CENP-A^ residences when compared to their ternary^Scm3^ counterparts (79 s vs. 108 s). Thus, physical restriction of the CEN DNA template, either by shortening it or limiting its mobility, prevents stable association of Cse4^CENP-A^, quantitatively reproducing the transient binding seen in the absence of either Ndc10 or the chaperone protein Scm3^HJURP^. These observations suggest that stable deposition of Cse4 requires physical wrapping of the CEN DNA around the histone core, that both Ndc10 and Scm3^HJURP^ have an important role in the wrapping process, and that wrapping of centromeric DNA is a key kinetic hurdle in CEN nucleosome formation.

**Table 1:**
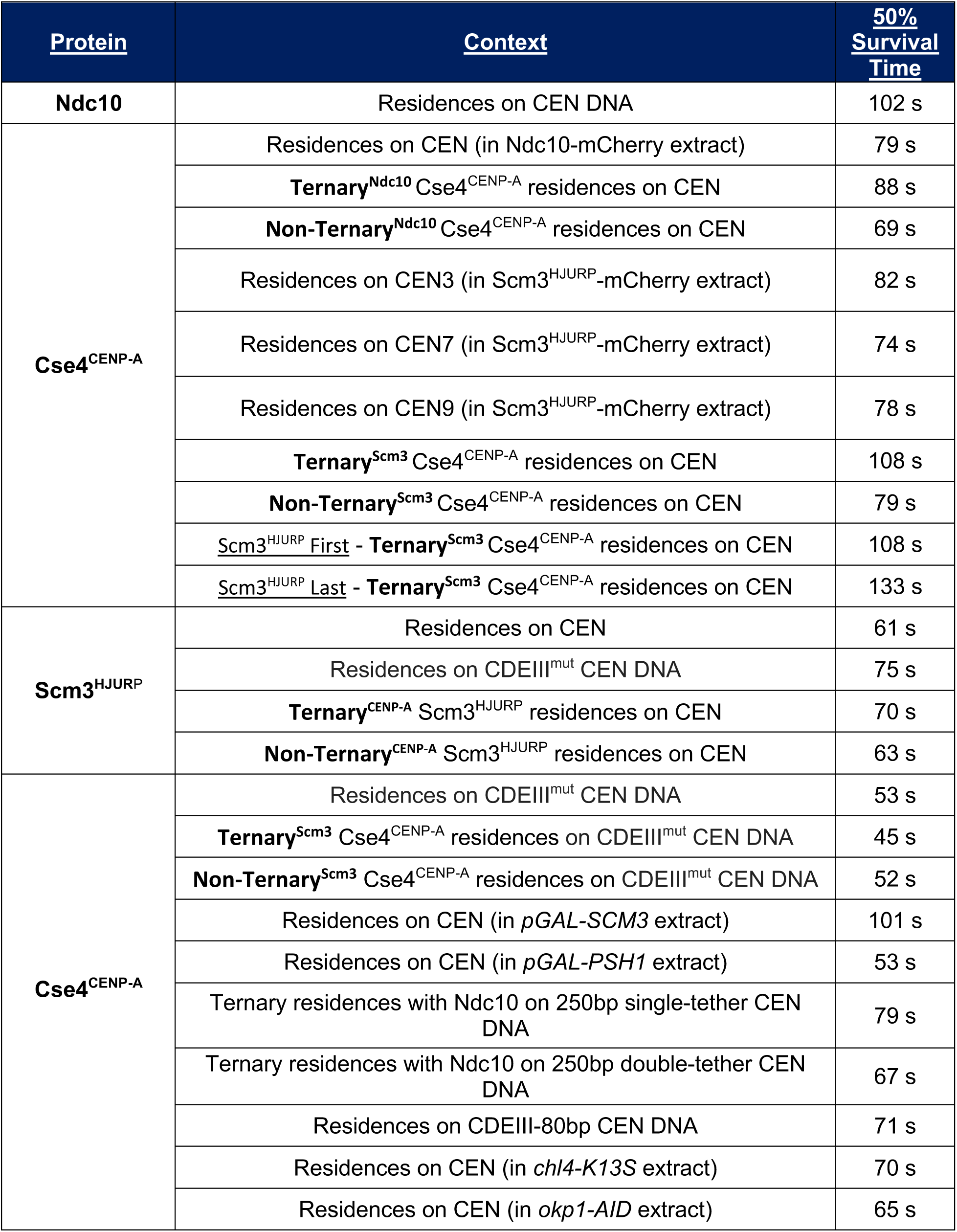
Median survival times of protein residences determined from various conditions tested and reported in this study.

| <u>Protein</u> | <u>Context</u> | <u>50% Survival Time</u> |
| --- | --- | --- |
| **Ndc10** | Residences on CEN DNA | 102 s |
| **Cse4<sup>CENP-A</sup>** | Residences on CEN (in Ndc10-mCherry extract) | 79 s |
|  | **Ternary<sup>Ndc10</sup> Cse4<sup>CENP-A</sup>** residences on CEN | 88 s |
|  | **Non-Ternary<sup>Ndc10</sup> Cse4<sup>CENP-A</sup>** residences on CEN | 69 s |
|  | Residences on CEN3 (in Scm3<sup>HJURP</sup>-mCherry extract) | 82 s |
|  | Residences on CEN7 (in Scm3<sup>HJURP</sup>-mCherry extract) | 74 s |
|  | Residences on CEN9 (in Scm3<sup>HJURP</sup>-mCherry extract) | 78 s |
|  | **Ternary<sup>Scm3</sup> Cse4<sup>CENP-A</sup>** residences on CEN | 108 s |
|  | **Non-Ternary<sup>Scm3</sup> Cse4<sup>CENP-A</sup>** residences on CEN | 79 s |
|  | <u>Scm3<sup>HJURP</sup> First</u> - **Ternary<sup>Scm3</sup> Cse4<sup>CENP-A</sup>** residences on CEN | 108 s |
|  | <u>Scm3<sup>HJURP</sup> Last</u> - **Ternary<sup>Scm3</sup> Cse4<sup>CENP-A</sup>** residences on CEN | 133 s |
| **Scm3<sup>HJURP</sup>** | Residences on CEN | 61 s |
|  | Residences on CDEIII<sup>mut</sup> CEN DNA | 75 s |
|  | **Ternary<sup>CENP-A</sup> Scm3<sup>HJURP</sup>** residences on CEN | 70 s |
|  | **Non-Ternary<sup>CENP-A</sup> Scm3<sup>HJURP</sup>** residences on CEN | 63 s |
| **Cse4<sup>CENP-A</sup>** | Residences on CDEIII<sup>mut</sup> CEN DNA | 53 s |
|  | **Ternary<sup>Scm3</sup> Cse4<sup>CENP-A</sup>** residences on CDEIII<sup>mut</sup> CEN DNA | 45 s |
|  | **Non-Ternary<sup>Scm3</sup> Cse4<sup>CENP-A</sup>** residences on CDEIII<sup>mut</sup> CEN DNA | 52 s |
|  | Residences on CEN (in *pGAL-SCM3* extract) | 101 s |
|  | Residences on CEN (in *pGAL-PSH1* extract) | 53 s |
|  | Ternary residences with Ndc10 on 250bp single-tether CEN DNA | 79 s |
|  | Ternary residences with Ndc10 on 250bp double-tether CEN DNA | 67 s |
|  | Residences on CDEIII-80bp CEN DNA | 71 s |
|  | Residences on CEN (in *chl4-K13S* extract) | 70 s |
|  | Residences on CEN (in *okp1-AID* extract) | 65 s |

### DNA-binding CCAN elements stabilize the centromeric nucleosome

Given the inherent instability of bare centromeric nucleosomes *in vitro* (Dechassa *et al*., 2014; Xiao *et al*., 2011), we hypothesized that additional stabilization after nucleosome formation might be provided by the binding of centromeric DNA-associated CCAN proteins. Such an additional stabilization step would explain our observations where Cse4^CENP-A^ and Scm3^HJURP^ colocalized together on CEN DNA without yielding long (>300 s) Cse4^CENP-A^ residences (Figure 3E). To test this idea, we focused on two conserved CCAN proteins that recent structural studies showed interact with CEN DNA, Chl4^CENP-N^ and Okp1^CENP-Q^ (Guan *et al*., 2021; Yan *et al*., 2019). Although CENP-C also exhibits significant DNA association within the reconstitution of the human CCAN (Yatskevich *et al*., 2022), we chose not to target Mif2^CENP-C^ due to its absence from the yeast structural model. Chl4^CENP-N^ binds close to the Cse4^CENP-A^ nucleosome and forms a DNA-binding groove that binds the centromere and stabilizes an extended DNA section adjacent to the nucleosome (Figure 5A). Mutation of this Chl4^CENP-N^ DNA-binding groove (*chl4^K13S^*) exhibits genetic interactions with mutants in Cse4^CENP-A^, making it an ideal candidate to test its contribution to nucleosome stability (Yan *et al*., 2019). Okp1^CENP-Q^ forms a heterodimer with Ame1^CENP-U^ that has DNA binding activity and has been proposed to interact with an N-terminal extension of Cse4^CENP-A^ within the nucleosome structure (Hinshaw & Harrison, 2019; Hornung *et al*, 2014; Yan *et al*., 2019) (Figure 5A). We therefore assayed Cse4^CENP-A^ endpoint colocalization in *chl4^K13S^*and *okp1-AID* extracts and found that it was significantly impaired, even though total Cse4^CENP-A^ levels in the extracts were not significantly altered (Figure 5B-D). Cse4^CENP-A^ residence lifetimes in both *chl4^K13S^* and *okp1-AID* extracts were also significantly reduced (Figure 5E, F). This disruption in centromeric nucleosome maintenance suggests that other kinetochore proteins that bind to CEN DNA also help to stabilize the nucleosome.

**Figure 5.**
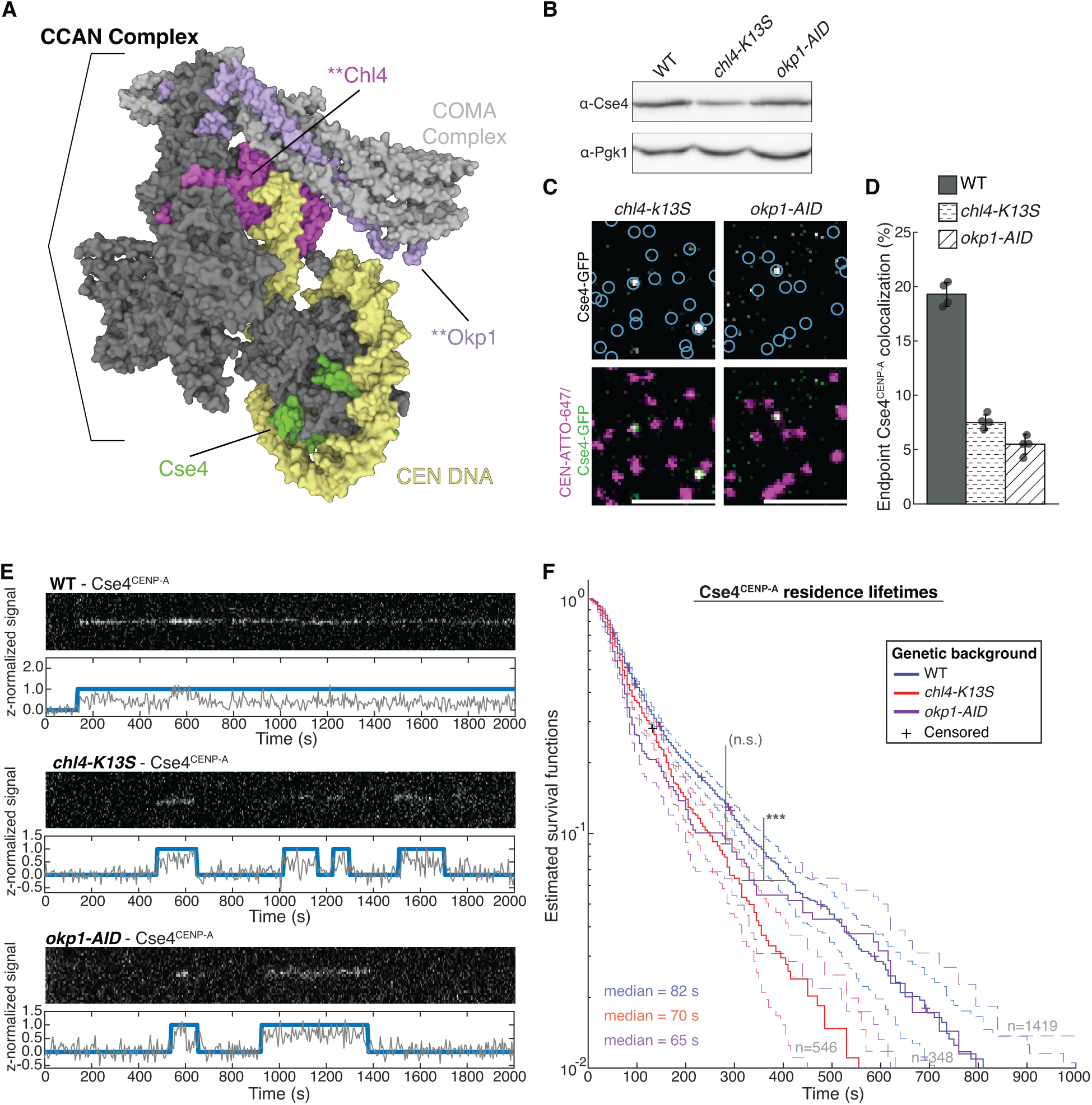
DNA-binding CCAN proteins stabilize the nucleosome to provide a platform for kinetochore assembly. A Structure of the yeast CCAN in complex with Cse4^CENP-A^ with CEN DNA (yellow), Cse4^CENP-A^ (green), Chl4^CENP-N^ (magenta) and Okp1^CENP-Q^ (purple), highlighting DNA adjacent regions targeted by the *chl4*-K13S mutant or proteasomal degradation of Okp1^CENP-Q^ (*okp1-AID)*. Image of 6QLD (Yan *et al*, 2019) created with Mol* (Sehnal *et al*, 2021). B Immunoblot analysis of whole cell extracts from WT, *chl4-K13S* and *okp1-AID* cells using indicated antibodies. C Example images of TIRFM endpoint colocalization assays. Top panels show visualized Cse4^CENP-A^-GFP on CEN DNA in extracts from *chl4*-*K13S* (top-left panel) or auxin-treated *okp1-AID* strains (*okp1-AID*, top-right panel) with colocalization shown in relation to identified CEN DNA in blue circles. Bottom panels show overlay of CEN DNA channel (magenta) with Cse4^CENP-A^-GFP (green), Scale bars 3 μm. D Quantification of Cse4^CENP-A^ endpoint colocalization with CEN DNA in extracts from WT, *chl4*-K13S, or *okp1-AID* genetic backgrounds (19 ± 1.1%, 8 ± 0.7%, 5 ± 0.9%, avg ± s.d. n=4 experiments, each examining ∼1,000 DNA molecules from different extracts). E Representative residence traces of Cse4^CENP-A^ signal on CEN DNA in WT (top), *chl4*-K13S (middle), or *okp1-AID* (bottom) extracts. Each example includes kymographs of Cse4^CENP-A^ (488 nm-top) with normalized intensity trace (grey-bottom) as well as identified residences (blue). Images acquired every 5 seconds with normalized fluorescence intensity shown in arbitrary units. F Kaplan-Meier analysis of Cse4^CENP-A^ residence lifetimes on CEN DNA in extracts from WT (blue median lifetime of 82 s, n=1419 over 3 experiments of ∼1000 DNA molecules using different extracts), *chl4-K13S* (red – median lifetime of 70 s, n=546 over 3 experiments of ∼1000 DNA molecules using different extracts) and *okp1-AID* (purple – median lifetime of 61 s, n=348 over 3 experiments of ∼1000 DNA molecules using different extracts) genetic backgrounds. Significant difference (***) between WT extract and *chl4*-K13S extract residence lifetime plots (two-tailed p-value of 3.4e-5 as determined by log-rank test). No significant difference (n.s.) between *chl4*-K13S and *okp1-AID* residence lifetimes in (two-tailed p-value of .40 as determined by log-rank test). 95% confidence intervals indicated (dashed lines), right-censored lifetimes (plus icons) were included and unweighted in survival function estimates.

### DNA composition of centromeres contributes to genetic stability through Cse4^CENP-A^ recruitment

We next sought to investigate the role of centromeric DNA sequence in centromeric nucleosome formation since it has been difficult to reconstitute stable centromeric nucleosomes in vitro (Guan et. al., 2021, Dechassa et. al., 2011, Yan et. al., 2019). To do so, we took advantage of earlier work, where a synthetic library of centromeric DNAs with randomized CDEII sequences was screened for members that can functionally replace native centromeres in vivo. This screen revealed that synthetic centromere function correlates strongly with high homopolymeric A + T content, defined as continuous “runs” of four or more A or T residues (A_n≥4_ or T_n≥4_), within CDEII (Baker & Rogers, 2005). Synthetic centromeres in which a high fraction of CDEII residues (> 0.38) occurred in runs of A_n≥4_ or T_n≥4_ were genetically more stable than synthetic centromeres with lower fractions of homopolymeric A + T runs within CDEII (< 0.38), although all chromosomes with synthetic centromeres were less stable in vivo when compared to a WT centromere. We therefore refer to the synthetic centromeres with more AT runs as “unstable” and those with less AT runs as “very unstable” mutants (Figure 6A). To test whether these differences in CDEII sequence and genetic stability correlate with the ability of the templates to retain Cse4^CENP-A^, we performed endpoint colocalization assays, using two unstable and two very unstable mutant CEN DNAs (Baker & Rogers). While all four mutants recruited lower levels of Cse4^CENP-A^ compared to WT CEN DNA, the unstable mutants recruited higher levels of Cse4^CENP-A^ than their very unstable counterparts (Figure 6B, C), and Cse4^CENP-A^ recruitment levels across the different templates correlated strongly with the fraction of CDEII residues in homopolymeric A + T runs (Figure 6D) and with the stability of chromosome inheritance in vivo (Figure 6D, inset). Time-lapse experiments with the same four mutants showed that the average number of Cse4^CENP-A^ residences observed per CEN DNA in a 45 min imaging sequence also correlated with homopolymeric A + T run content and CEN functionality in vivo (Figure 6E, left). This relationship was apparently independent of Scm3^HJURP^, because the average number of Scm3^HJURP^ residences observed per CEN DNA remained consistent across the mutants (Figure 6E, right). Consistent with this Scm3^HJURP^ independent behavior, the proportion of ternary Cse4^CENP-A^-Scm3^HJURP^ residences was similar in “Very Unstable” CDEII mutant CEN DNA when compared to CEN DNA (Appendix Figure S3A), but there was a significant increase in the estimated off-rates of ternary^Scm3^ Cse4^CENP-A^ residences on “Very Unstable” CDEII mutant CEN DNA (Appendix Figure S3B). These results suggest that despite not affecting Scm3^HJURP^-Cse4^CENP-A^ ternary residence initiation, very unstable CDEII mutants might restrict Scm3^HJURP^ catalyzed stabilization of Cse4^CENP-A^.

**Figure 6.**
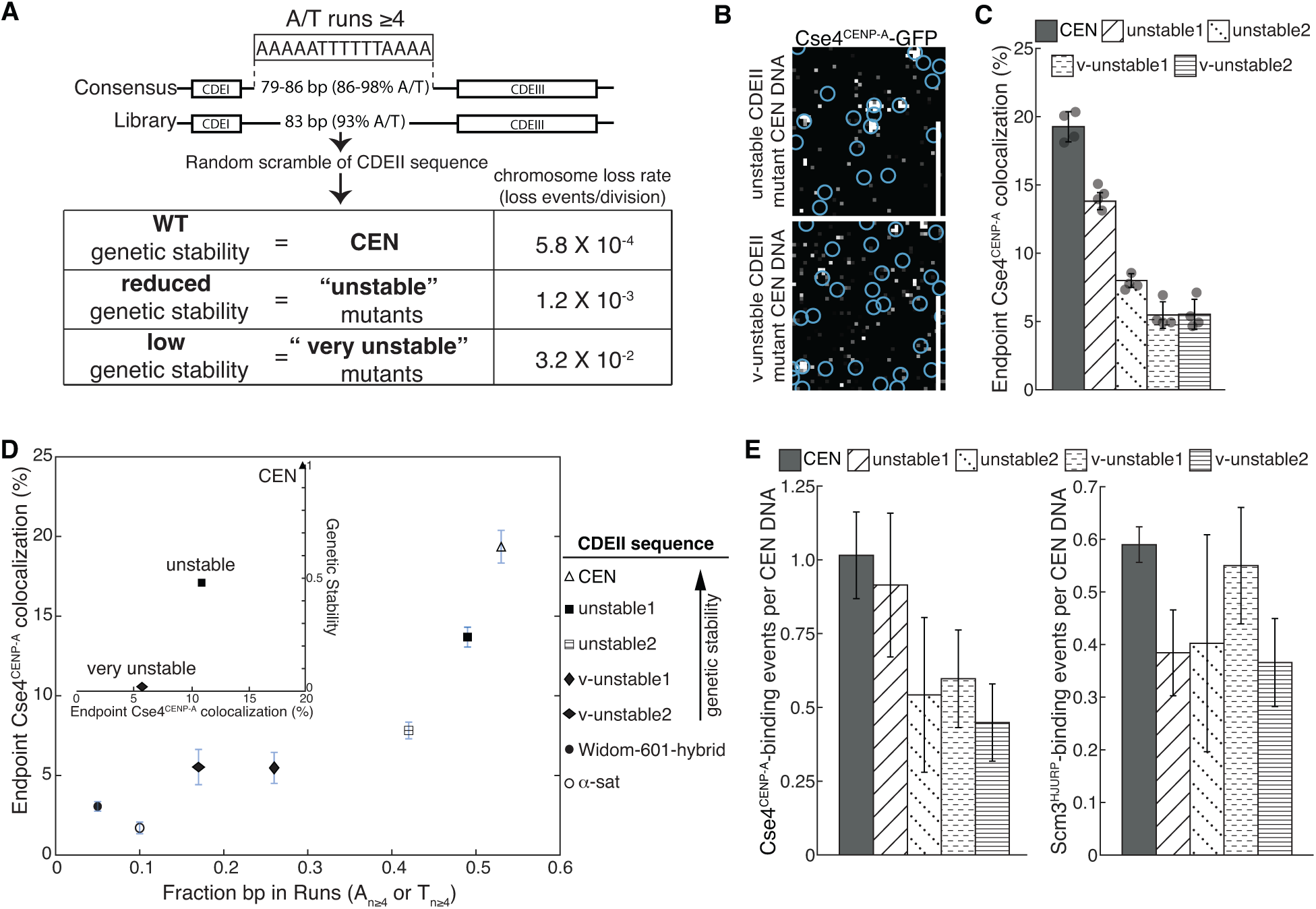
DNA-composition of centromeres contributes to genetic stability through Cse4^CENP-A^ recruitment. A Overview of CDEII mutants generated for stability assays where overall % of A/T content was maintained while A/T run content was randomly varied and selected for genetic stability including reported chromosome loss rates of WT and CDEII mutant pools (adapted from (Baker & Rogers, 2005). B Example images of TIRFM endpoint colocalization assays. Visualized Cse4^CENP-A^-GFP on unstable1 CDEII-mutant DNA (top panel) or on v-unstable1 CDEII-mutant DNA (bottom panel) with colocalization shown in relation to identified CEN DNA in blue circles. Scale bars 3 μm. C Quantification of endpoint colocalization of Cse4^CENP-A^ on CEN, unstable1, unstable2, v-unstable1, and v-unstable2 CEN DNA (19.3 ± 1.1%, 13.8 ± 0.6%, 7.8 ± 0.5%, 5.5 ± 1.0%, 5.5 ± 1.1%, avg ± s.d. n=4 experiments, each examining ∼1,000 DNA molecules from different extracts). D Stable Cse4^CENP-A^ recruitment depends upon CDEII sequence A/T run content. Plot of fraction of all CDEII bp that occur in homopolymeric A_n≥4_ or T_n≥4_ repeats (fraction bp in runs (N≥4) in CEN, unstable1, unstable2, v-unstable1, v-unstable2, Widom-601 hybrid and α-sat CEN DNA (0.53, 0.49, 0.41, 0.26, 0.17, 0.05, 0 .10), versus the observed colocalization of Cse4^CENP-A^ on CEN, unstable1 unstable2, v-unstable1, v-unstable2, Widom-601 hybrid and α-sat CEN DNA (19.3 ± 1.1%, 13.8 ± 0.6%, 7.8 ± 0.5%, 5.5 ± 1.0%, 5.5 ± 1.1%, 3.1 ± 0.3%, 1.7 ± 0.4% (avg ± s.d. n=4). Inset plot of Cse4^CENP-A^ endpoint colocalization percentage on CEN, unstable mutants (average), and very unstable mutants (average) (19.9%, 10.8% and 5.5% respectively) versus genetic stability (chromosome loss normalized to CEN) of WT, unstable mutants, and very unstable mutants (1.0, 0.48 and 0.02 respectively). E Very unstable CDEII mutants have reduced average Cse4^CENP-A^ binding when compared to unstable counterparts. Average residences of Cse4^CENP-A^ per CEN DNA (left) on CEN, unstable1, unstable2, v-unstable1, and v-unstable2 CEN DNA (1.02 ± 0.15, 0.92 ± 0.24, 0.54 ± 0.26, 0.60 ± 0.17, 0.45 ± 0.13, avg ± s.e.m. n=3 experiments of ∼1000 DNA molecules using different extracts) and average residences of Scm3^HJURP^ per CEN DNA (right) on CEN, unstable1, unstable2, v-unstable1, and v-unstable2 CEN DNA (0.59 ± 0.03, 0.38 ± 0.08, 0.40 ± 0.21, 0.55 ± 0.11, 0.37 ± 0.08, avg ± s.e.m. n=3 experiments of ∼1000 DNA molecules using different extracts).

To further explore the role of DNA sequence, we generated several other templates that maintained high percentages of A and T residues (high % A/T) yet had very little homopolymeric A_n≥4_ or T_n≥4_ run content. A CDEII mutant that contained a substitution of satellite DNA from human chromosomes (α-sat) failed to recruit stable Cse4^CENP-A^ in endpoint colocalization assays despite its high % A/T (Figure 6D). The hybrid sequence combining CDEIII and Widom 601 (Widom-601-hybrid) that readily forms centromeric nucleosomes in recombinant reconstitutions (Yan *et al*., 2019) (Appendix Figure S4A) also failed to recruit stable Cse4^CENP-A^ in endpoint colocalization assays (Figure 6D). To ensure that the Widom-601-hybrid CEN DNA was not saturated by the canonical H3 histone, we also monitored H3 incorporation. The Widom-601-hybrid CEN DNA incorporated low levels of H3, similar to WT CEN DNA, in both bulk assembly assays (Appendix Figure S4B) and endpoint colocalization assays (Appendix Figure S4C, D). Such low levels of H3 incorporation were unexpected, as Widom-601 readily forms stable H3 nucleosomes when reconstituted (Lowary & Widom, 1998), but this may point to differences in *de novo* histone formation that have yet to be fully studied. Removal of the CDEIII sequence from the Widom-601-hybrid to prevent CBF3 binding yielded an approximately 2-fold increase in H3 incorporation (Appendix Figure S4C, D), suggesting that CBF3 may negatively regulate H3 binding at centromeres. Taken together, these data highlight a critical functional role for homopolymeric A + T run content within CDEII for stable Cse4^CENP-A^ recruitment.

## Discussion

Here, we report adaptation of a cell-free system to autonomously assemble native centromeric nucleosomes on individual centromeric DNA molecules with CoSMoS to enable the study of native centromeric nucleosome formation at spatiotemporal resolutions not previously accessible. Continuous monitoring revealed that several cofactors coordinate to promote stable Cse4^CENP-A^ recruitment and maintenance at the centromere. These cofactors include the DNA-binding CBF3 component Ndc10, as well as the conserved chaperone Scm3^HJURP^. We found that Scm3^HJURP^ was a limiting cofactor that promoted stable centromeric Cse4^CENP-A^ association but was not required for transient centromere association. Stabilization of Cse4^CENP-A^ likely occurred through catalysis of centromeric DNA wrapping because we observed that physical restriction of the DNA impaired stable recruitment of Cse4^CENP-A^. Guided by recent structural kinetochore reconstitutions (Yan et. al., 2019, Guan et. al., 2021), we also found that the centromeric nucleosome must be stabilized by DNA-associated kinetochore proteins within the CCAN, highlighting the tight coordination of kinetochore assembly at the inner kinetochore. This finding may in part explain the instances where Cse4^CENP-A^ colocalized on CEN DNA with its chaperone Scm3^HJURP^ yet failed to remain stably associated. In those instances, the Cse4^CENP-A^ nucleosome may have successfully formed, but subsequent association of CCAN kinetochore proteins failed to occur, permitting dissolution of the nucleosome complex (Figure 7). We also interrogated the CDEII DNA element and identified a role for sequence composition in stable centromeric nucleosome recruitment which correlated with centromere stability in cells. Taken together, this assay enabled the assessment at high resolution of the native nucleosome assembly process on centromeric DNA as well as a functional role of centromere DNA sequence composition.

**Figure 7.**
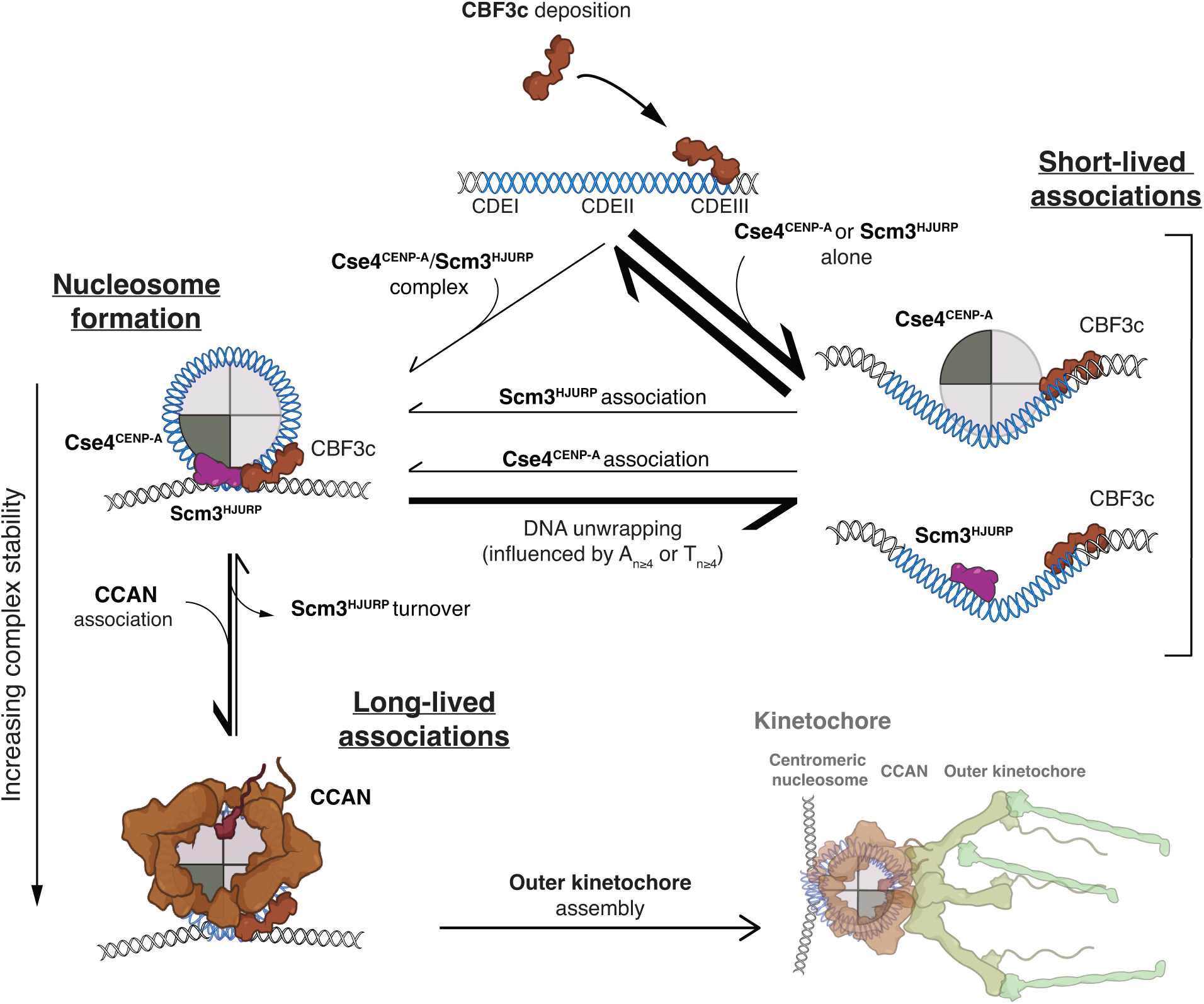
Formation of stable centromeric nucleosome requires tight coordination of centromeric DNA, Cse4^CENP-A^ with its chaperone Scm3^HJURP^ and CCAN kinetochore protein association. Schematic of centromeric nucleosome formation highlighting the different pathways that could lead to either short-lived or long-lived residences Cse4^CENP-A^ as measured in TIRFM resident lifetime assays.

### Formation of a native centromeric nucleosome

Despite over a decade of ongoing study, reconstituting Cse4^CENP-A^ nucleosomes with yeast centromeric DNA has remained remarkably elusive. Recent structural studies have started to shed light on these complexes, yet there are fundamental differences between these reconstitutions and native kinetochore assemblies. One such example is our finding that the Widom 601-hybrid sequence that readily stabilized the Cse4^CENP-A^ nucleosome *in vitro* was a poor template for Cse4^CENP-A^ recruitment and stable nucleosome formation *de novo* (Figure 6E). Structural models using non-native CEN DNA required significant rearrangements around the centromeric nucleosome to permit CCAN assembly, including dissociation of the CBF3 complex (Guan *et al*., 2021; Yan *et al*., 2019). However, we did not observe a significant reduction in CBF3 complex colocalization, even after sufficient incubation to allow full *de novo* kinetochore assembly (Lang *et al*., 2018). Our findings are consistent with observed CBF3 behavior in cells (Joglekar *et al*., 2008) and highlight both the potential significant differences between *in vitro* reconstitutions of kinetochore complexes and their native counterparts as well as the significant advantage of assembling kinetochores under native conditions.

Our work suggests the major functional role of the Scm3^HJURP^ chaperone at centromeres may be Cse4^CENP-A^ incorporation into stable nucleosomes instead of centromere targeting (Camahort *et al*., 2007; Cho & Harrison, 2011b). We found that Scm3^HJURP^ is a limiting kinetochore assembly factor required to catalyze stable Cse4^CENP-A^ recruitment (Figure 3, 4). The population of Cse4^CENP-A^ that localized to CEN DNA in the absence of Scm3^HJURP^ was presumably bound to other chaperone proteins it is known to associate with, such as CAF-1, Spt6, and DAXX (Bobkov *et al*, 2020; Hewawasam *et al*, 2018; Lacoste *et al*, 2014). It should be acknowledged that it is possible that transient Cse4^CENP-A^ binding may be driven in some part through mass action due to high concentrations of CEN DNA and *in vivo*, this transient binding may not be physiologically relevant. However, our data shows that these associations are insufficient for stable nucleosome formation in the absence of Scm3^HJURP^. Given that physical restriction of DNA wrapping severely limited stable association of Cse4^CENP-A^ (Figure 4), we further propose that Scm3^HJURP^ promotes stable Cse4^CENP-A^ recruitment through catalysis of centromeric DNA wrapping, consistent with its activity in vitro (Dechassa et. al., 2011, Xiao et.al., 2011). While several chaperones have been shown to limit ectopic Cse4^CENP-A^ deposition (Gkikopoulos *et al*, 2011; Hewawasam *et al*., 2018; Ranjitkar, 2010), it is likely that Scm3^HJURP^ promotes specific deposition at centromeres through two non-exclusive mechanisms: (1) tethering of Cse4^CENP-A^ via Scm3^HJURP^-Ndc10 binding to promote centromeric DNA wrapping and (2) stabilization of a Cse4^CENP-A^ centromere DNA intermediate through the AT-rich DNA binding domain of Scm3^HJURP^ (Xiao *et al*., 2011). While we aim to further investigate this mechanism, either possibility yields a model of Scm3^HJURP^-catalyzed Cse4^CENP-A^ nucleosome formation that is supported by *in vitro* reconstitutions, where it was found that Scm3^HJURP^ was required to form a Cse4^CENP-A^ nucleosome on the highly AT-rich (and thus inherently unfavorably for histone wrapping) centromeric DNA (Dechassa *et al*., 2011; Xiao *et al*., 2011). In addition, this model is consistent with *in vivo* studies that showed Scm3^HJURP^ coordinates with Ndc10 to deposit Cse4^CENP-A^ at the centromere and is persistent at centromeres after Cse4^CENP-A^ deposition, although undergoing rapid exchange (Wisniewski *et al*., 2014; Xiao *et al*., 2011).

### A functional role for the essential centromere element CDEII

The single molecule TIRFM colocalization assays developed here enabled direct assessment of a functional role of the centromeric CDEII element, a question that has been extremely difficult to address *in vivo*. While there is no sequence conservation across CDEII elements in yeast, they are often among the loci with the highest A/T % across the genome (Baker & Rogers, 2005). Although high A/T % is not a universal feature of all centromeres (Melters *et al*, 2013), it is a widely conserved feature across centromeres in both yeast and higher organisms despite being unfavorable for core histone wrapping (Struhl & Segal, 2013). Our work uncovered a role for CDEII sequence composition in stable Cse4^CENP-A^ nucleosome recruitment and suggests that homopolymeric A + T run content contributes to stable incorporation of Cse4^CENP-A^. The requirement for A + T run content is highlighted by the failure of the canonical nucleosome-targeting Widom 601-based centromere (Widom 601-hybrid), which has high A/T % but contains little A + T run content, to stably recruit Cse4^CENP-A^ in our assay (Figure 6E), despite readily forming in reconstitutions (Xiao *et al*., 2011). This points to potentially significant differences in the formation of native centromeric complexes and their recombinant counterparts. In addition, it remains an open question whether yeast CDEII element sequences have evolved to maximize Cse4^CENP-A^ recruitment and stability or have evolved to a point where Cse4^CENP-A^ stability is sufficient to maintain centromere identity, as no A + T run beyond 8 nucleotides occurs within CDEII sequences despite occurring outside of centromeric loci (Baker & Rogers, 2005). Organisms with more complex centromeric architectures that maintain high A/T% fail to substitute for CDEII function in yeast centromeres, likely due to the lack of high A + T run content (Figure 6E).

Our work also supports a previously proposed model where A/T-rich centromeric DNA functions to restrict canonical histone formation (Dechassa *et al*., 2014; Drew & Travers, 1985; Stormberg & Lyubchenko, 2022; Xiao *et al*., 2011). This model is consistent with findings in other organisms where H3 histone eviction was an inherent property of centromeric DNA and with our own findings that H3 has low occupancy on the CEN templates (Baker & Rogers, 2005; Shukla *et al*., 2018). This potential functional role of centromere sequence and/or centromere-binding kinetochore proteins in H3-eviction may be providing a drive to the genetic conservation of AT-rich DNA amongst centromere sequences. We speculate that in yeast, additional DNA-binding kinetochore proteins arose to aid in overcoming this kinetic barrier and to protect centromere function. This may be a consequence of the fact that despite a lack of meiotic drive, yeast centromeres are among the fastest evolving regions of the yeast genome and are thus sensitive to negative selection resulting in complete loss of centromere identity (Bensasson *et al*, 2008). Rapid purifying selection may also explain the accumulation of homopolymeric A+ T run content within CDEII, which we found correlated to stable Cse4^CENP-A^ recruitment. This was somewhat unexpected, as homopolymeric repeats were found to resist nucleosome formation (Liebl & Zacharias, 2021). However, it has been reported that stretches of A-repeats impose a net curvature into dsDNA and that the DNA bend at centromeres contributes to their stability in cells (Beutel & Gold, 1992; Murphy *et al*, 1991; Widom, 2001). We therefore propose that homopolymeric A + T runs are cooperative with additional DNA-associated kinetochore proteins to facilitate nucleosome stability and kinetochore assembly and may explain the observed requirement for the A/T-rich DNA binding Scm3^HJURP^ to catalyze centromeric DNA wrapping of Cse4^CENP-A^ (Figure 3, 4). The precise mechanism by which these A + T runs contribute to nucleosome formation remains open for further study, but at the very least may provide context for the required coordination of several DNA-binding proteins that are in close proximity to the Cse4^CENP-A^ nucleosome (Figure 7).

Adaptation of the assay developed here to study downstream kinetochore assembly will enable the study of the contributions of various kinetochore proteins from within the vast network required to form a functional kinetochore scaffold. This assay may also be adapted to study the dynamics of other histones and chaperones as well as to further recapitulate a native centromere “cell-like” environment by including processes such as centromere replication, which precludes kinetochore assembly in cells (Bobkov *et al*., 2020; Furuyama & Biggins, 2007; Gkikopoulos *et al*., 2011). In addition, templates that include more complex chromatin structures surrounding the centromere that may provide additional information about the roles pericentromeric chromatin plays in maintenance of centromere identity and function. Such adaptations may be needed to better understand how such an initially tenuous assembly pathway with stringent prerequisite conditions occurs so rapidly and with such fidelity in cells throughout passage of every cell cycle. The mechanisms that drive this process and that may be abrogated in various cellular disease states are a critical area of study.

## Materials and Methods

### Yeast Methods

The *S. cerevisiae* strains used in this study are listed in Supplemental Table 1 and are derivative of SBY3 (W303). Standard genetic crosses, media and microbial techniques were used. Cse4 was tagged internally at residue 80 with eGFP including linkers on either side (pSB1617) and then expressed from its native promoter at an exogenous locus in a *cse4#x03B4;* background (SBY19926). Genes that were changed to include endogenously tagged fluorescent protein alleles (mCherry) or epitope tags (- 13myc) or auxin-inducible degrons (-IAA7) were constructed at the endogenous loci by standard PCR-based integration techniques (Longtine *et al*, 1998) and confirmed by PCR. The mutant *chl4-K13S* was made via PCR-based integration from a vector (pSB2182) containing the *CHL4* gene with 13 mutations present (Yan *et al*., 2019). The plasmids and primers used to generate strains are listed in Supplemental Tables 2 and 3, respectively. All liquid cultures were grown in yeast peptone dextrose rich (YPD) media. To arrest cells in mitosis, log phase cultures were diluted in liquid media to a final concentration of 30 μg/mL benomyl and grown for another three hours until at least 90% of cells were large budded. For strains with auxin inducible degron (AID) alleles (*scm3-AID*, *okp1-AID*), all cultures used in the experiment were treated with 500 μM indole-3-acetic acid (IAA, dissolved in DMSO) for the final 60 min of growth as described previously (Lang *et al*., 2018; Miller *et al*, 2016; Nishimura *et al*., 2009). For strains with galactose inducible alleles (*pGAL-PSH1*, *pGAL-SCM3*), cultures were grown in raffinose and then treated with 4% galactose for the final 60 min of growth. Growth assays were performed by diluting log phase cultures to OD600 ∼ 1.0 from which a 1:5 serial dilution series was made. This series was plated on YPD and YP plates that contained 4% Galactose and incubated at 23 °C.

### Preparation of DNA templates and Dynabeads

Plasmid pSB963 was used to generate the WT CEN DNA templates and pSB972 was used to generate the CEN^mut^ template used in this study. Derivatives of pSB963 were altered using mutagenic primers and Q5 site-directed mutagenesis kit (NEB) to generate vectors containing CEN9, unstable1, unstable2, very unstable1, very unstable2, α-sat, and 80bp-CDEIII sequences (pSB3338, pSB3336, pSB3416, pSB3337, pSB3415 and pSB3335, respectively). CDEII mutants were taken directly from high-loss and low-loss pools from Baker & Rodgers, 2005, where very unstable1 is equivalent to H1, very unstable2 is equivalent to H14, unstable1 is equivalent to L1, and unstable2 is equivalent to L5. Widom-601 template was equivalent to the one used in structural studies (Guan *et al*., 2021) where CENIII (92-137) was inserted into corresponding region of 601 sequence (pSB3264). For α-sat CEN DNA, the CDEII region of the CEN template was mutated to a fragment of α-satellite DNA from *H. sapiens* X-chromosome based on structural studies containing CENP-A (Yatskevich *et al*., 2022). DNA templates were generated by PCR using a 5’-ATTO647-funtionalized (IDT DNA) 5’ primer with homology to linker DNA upstream of ∼60 bp of pericentromeric DNA and the centromere (SB7843) and a 5’-biotinylated (IDT DNA) 3’ primer with linker DNA, an EcoRI restriction site, and homology to linker DNA downstream of ∼ 60 bp of pericentromeric DNA and the centromere (SB3879) to yield ∼750 bp dye-labeled assembly templates. For the 80 bp CDEIII mutant, plasmid pSB963 was amplified with primers SB7843 and SB7844 to generate a CDEIII containing mutant of 80 bp total length. For CEN7, template pSB2953 was amplified with primers SB5699 and SB7842 to generate a dye-labeled and biotinylated 750 bp assembly template. For the double-tethered CEN DNA template, biotinylated primers SB7845 and SB3878 were used to generate a 250 bp template. The single-tethered 250 bp control template was generated using primers SB7846 and SB3878. The Widom-601 template (pSB2887) and Widom-601-CDEIII hybrid template (pSB3264) were amplified with primers SB5699 and SB7842 to generate dye-labeled and biotinylated 750 bp templates. Supplemental Table 2 includes the plasmids used in this study and Supplemental Table 3 includes the primer sequences used to PCR amplify the DNA templates. PCR products were purified using the Qiagen PCR Purification Kit. In the case of bulk assembly for Mnase assays, purified CEN DNA was conjugated to Streptadivin-coated Dynabeads (M-280 Streptavidin, Invitrogen) for 2.5 hr at room temperature, using 1 M NaCl, 5 mM Tris HCl (pH7.5), and 0.5 mM EDTA as the binding and washing buffer. For single molecule TIRFM assays, purified DNA was diluted in dH2O to a final concentration ∼100pM.

### Whole Cell Extract preparation for kinetochore assembly assays

For a standard bulk kinetochore assembly assay *in vitro*, cells were grown in liquid YPD media to log phase and arrested in mitosis in 500 mL and then harvested by centrifugation. All subsequent steps were performed on ice with 4 °C buffers. Cells were washed once with dH2O with 0.2 mM PMSF, then once with Buffer L (25 mM HEPES pH 7.6, 2 mM MgCl2, 0.1 mM EDTA pH 7.6, 0.5 mM EGTA pH 7.6, 0.1 % NP-40, 175 mM K-Glutamate, and 15% Glycerol) supplemented with protease inhibitors (10 mg/ml leupeptin, 10mg/ml pepstatin, 10mg/ml chymostatin, 0.2 mM PMSF), and 2 mM DTT. Cell pellets were then snap frozen in liquid nitrogen and then lysed using a Freezer/Mill (SPEX SamplePrep), using 10 rounds that consisted of 2 min of bombarding the pellet at 10 cycles per second, then cooling for 2 min. The subsequent powder was weighed and then resuspended in Buffer L according to the following calculation: weight of pellet (g) x 2=number of mL of Buffer L. Resuspended cell lysate was thawed on ice and clarified by centrifugation at 16,100 g for 30 min at 4 °C, the protein-containing layer was extracted with a syringe and then aliquoted and snap frozen in liquid nitrogen. The resulting soluble whole cell extracts (WCE) generally had a concentration of 50–70 mg/mL. The pellets, powder, and WCE were stored at -80 °C.

### Bulk assembly assays followed by Mnase treatment

De novo kinetochore assembly was performed with whole cell extract from SBY21110 (Ndc10-mCherry, Cse4-GFP) as previously described (Lang *et al*., 2018). Briefly, 1 mL of whole cell extract and 50 μl of DNA coated M280 Dynabeads (single-tether CEN, double tether CEN, or CEN^mut^) were incubated at room temperature for 90 min to allow kinetochore assembly. Following the final wash, beads were resuspended in 90 μL of Buffer L (see above) supplemented with 100 μg/mL BSA, 10 μL of 10X Micrococcal Nuclease Reaction Buffer (NEB) and 1000 gel units of Micrococcal Nuclease (NEB, #M0247S) and incubated at 30 °C for 10 min under constant mixing. The reaction was stopped with addition of EGTA to 10 mM. After removal of magnetic beads, the aqueous phase was phenol-extracted followed by ethanol precipitation of DNA and resuspended in dH20 and run on a 1% agarose gel. Resolved DNA were visualized via SYBR Gold nucleic acid dye (Invitrogen; S11494) on ChemiDoc Imager (Bio-Rad).

### Immunoblotting

For immunoblots, proteins were transferred from SDS-PAGE gels onto 0.22 μM cellulose paper, blocked at room temperature with 4% milk in PBST, and incubated overnight at 4°C in primary antibody. Antibody origins and dilutions in PBST were as follows: α-Cse4 (9536 (Pinsky *et al*, 2003); 1:500), α-H3 Alexa Fluor 555 (Invitrogen; 17H2L9; 1:3,000), α-PGK1 (Invitrogen; 4592560; 1:10,000). The anti-Scm3 antibodies were generated in rabbits against a recombinant Scm3 protein fragment (residues 1-28) of the protein by Genscript. The company provided affinity-purified antibodies that we validated by immunoprecipitating Scm3 from yeast strains with Scm3-V5 and confirming that the antibody recognized a protein of the correct molecular weight that was also recognized by α-V5 antibody (Invitrogen; R96025; 1:5000). We subsequently used the antibody at a dilution of 1:10,000. Secondary antibodies were validated by the same methods as the primary antibodies as well as with negative controls lacking primary antibodies to confirm specificity. Blots were then washed again with PBST and incubated with secondary antibody at room temperature. Secondary antibodies were α-mouse (NA931) or α-rabbit (NA934), horseradish peroxidase-conjugated purchased from GE Healthcare and used at 1:1000 dilution in 4% milk in PBST. Blots were then washed again with PBST and ECL substrate from Thermo Scientific used to visualize the proteins on ChemiDoc Imager (Bio-Rad). Uncropped and unprocessed scans of gels and blots are provided in the Source Data file.

### Single molecule TIRFM slide preparation

Coverslips and microscope slides were ultrasonically cleaned and passivated with PEG as described previously (Larson & Hoskins, 2017; Larson *et al*, 2014). Briefly, ultrasonically cleaned slides were treated with vectabond (Vector Laboratories) prior to incubation with 1% (w/v%) biotinylated mPEG-SVA MW-5000K/mPEG-SVA MW-5000K (Lysan Bio) in flow chambers made with double-sided tape. Passivation was carried out overnight at 4 °C. After passivation, flow chambers were washed with Buffer L and then incubated with 0.3 M BSA/0.3M Kappa Casein in Buffer L for 5 min. Flow chambers were washed with Buffer L and then incubated with 0.3M Avidin DN (Vector Laboratories) for 5 min. Flow chambers were then washed with Buffer L and incubated with ∼100 pM CEN DNA template for 5 min and washed with Buffer L. For endpoint colocalization assays, slides were prepared as follows: Flow chambers were filled with 100 μL of WCE containing protein(s) of interest via pipetting and wicking with filter paper. After addition of WCE, slides were incubated for 90 min at 25°C and then WCE was washed away with Buffer L. Flow chambers were then filled with Buffer L with oxygen scavenger system (Aitken *et al*, 2008) (10 nM PCD/2.5 mM PCA/1mM Trolox) for imaging. For immunofluorescence of H3, after 90 min, chambers were washed with Buffer L and then incubated for 30 min with Buffer L and 1:300 diluted antibody (Invitrogen 17H2L9). Chambers were then washed with Buffer L prior to imaging in Buffer L with oxygen scavenger system (above). For real-time colocalization assays, slides were prepared as follows: On the microscope, 100 μL WCE spiked with oxygen scavenger system was added to flow chamber via pipetting followed by immediate image acquisition.

### Single molecule TIRFM colocalization assays image collection and analysis

All images were collected on a Nikon TE-2000 inverted TIRF microscope with a 100x oil immersion objective (Nikon Instruments) with an Andor iXon X3 DU-897 EMCCD camera. Images were acquired at 512 px x 512 px with a pixel size of .11 µm/px at 10MHz. Atto 647 labeled CEN DNAs were excited at 640 nm for 300 ms, GFP-tagged proteins were excited at 488 nm for 200 ms, and mCherry-tagged proteins were excited at 561 nm for 200 ms. For endpoint colocalization assays, single snapshots of all channels were acquired. For real-time colocalization assays images in 561 nm channel and 488 nm channel were acquired every 5 s with acquisition of the DNA-channel (647 nm) every 1 min for 45 min total (541 frames) using Nikon Elements acquisition software. Snapshots were processed in a CellProfiler 4 image analysis (Stirling *et al*, 2021) pipeline using RelateObjects module to determine colocalization between DNA channel (647 nm) and GFP (488 nm) and mCherry (561 nm) channels. Results were quantified and plotted using MATLAB (The Mathworks, Natick, MA). Adjustments to example images (contrast, false color, etc.) were made using FIJI (Schindelin *et al*, 2012) and applied equally across entire field of view of each image.

For real-time colocalization assay, a custom-built image analysis pipeline was built in MATLAB (R2019b) to extract DNA-bound intensity traces for the different fluorescent species, to convert them into ON/OFF pulses and to generate the empirical survivor function data. First, the image dataset was drift-corrected using either fast Fourier Transform cross-correlations or translation affine transformation depending on the severity of the drift. DNA spots were identified after binarizing the DNA signal using global background value as threshold, as well as size and time-persistency filtering. Mean values of z-normalized fluorescent markers intensities were measured at each DNA spot at each time frame, and local background was subtracted. Z-normalized traces were then binarized to ON/OFF pulses by applying a channel-specific, manually adjusted threshold value unique to all traces in a given image set. Pulse onsets, durations and overlaps between channels were then derived. For plotting clarity, z-normalized traces shown in the figures were zero-adjusted so that the baseline signal lies around zero. Pulses in ON state at the end of the image acquisition (right censored) were annotated and included as unweighted in lifetime estimates. Kaplan-Meier analysis and log rank tests were performed in MATLAB (R2021a). Adjustments to example plot images (contrast) as well as generation of example plot source movies were made using FIJI (Schindelin *et al*., 2012).

### Statistical tests

Right-censored lifetimes were included in an unweighted Kaplan-Meier analysis to estimate survival functions, censored events typically comprised <5% of observed lifetimes. Survival functions including 95% confidence intervals were plotted using KMplot (Curve Cardillo G. (2008). KMPLOT: Kaplan-Meier estimation of the survival function. http://www.mathworks.com/matlabcentral/fileexchange/22293) and P-values for comparing the Kaplan–Meier survival plots are provided in the figure legends and were computed using the log-rank test within Logrank (Cardillo G. (2008). LogRank: Comparing survival curves of two groups using the log rank test http://www.mathworks.com/matlabcentral/fileexchange/22317). Different lifetime plots were considered significantly different with a *P-*value less than 0.05. Significance between off-rates were determined by two-tailed unpaired t-tests.

### Data Availability Section

Custom software written in MATLAB (R2021a) was used for TIRF colocalization residence lifetime analysis and plot generation. The source code is publicly available at https://github.com/FredHutch/Automated-Single-Molecule-Colocalization-Analysis. All primary datasets used for analysis in all figures are available at Zenodo, https://zenodo.org/ and assigned the identifier (10.5281/zenodo.7939301). All primary data reported in this paper will be shared by the lead contact upon request.

## Acknowledgements

All imaging was performed at the Fred Hutchinson Cancer Center Cellular Imaging Core (supported by the Fred Hutch/University of Washington Cancer Consortium P30 CA015704), and we thank Jin Meng and Lena Schroeder for their experimental help. We also thank members of the S.B. and C.L.A. labs for critical reading of the manuscript.

A.R.P. was supported by postdoctoral fellowship NIH F32GM136010. J.D.L was supported by CVP training grant 5T32HL7312-39 and a Washington Research Foundation Fellowship Award. C.L.A. was supported by R01GM079373 and R35GM134842. S.B. was supported by NIH R01GM064386 and is also an investigator of the Howard Hughes Medical Institute.

## Author contributions

A.R.P conceptually designed and performed experiments, analyzed data, and wrote the manuscript with input from all authors; J.D.L. conceptually designed experiments and edited the manuscript. J.D. designed data analysis software and reviewed and edited the manuscript. C.L.A. and S.B. conceptually designed and supervised experiments and reviewed and edited the manuscript.

## Disclosure and competing interests statement

The authors declare no conflict of interest.

---

**Figure EV1.**
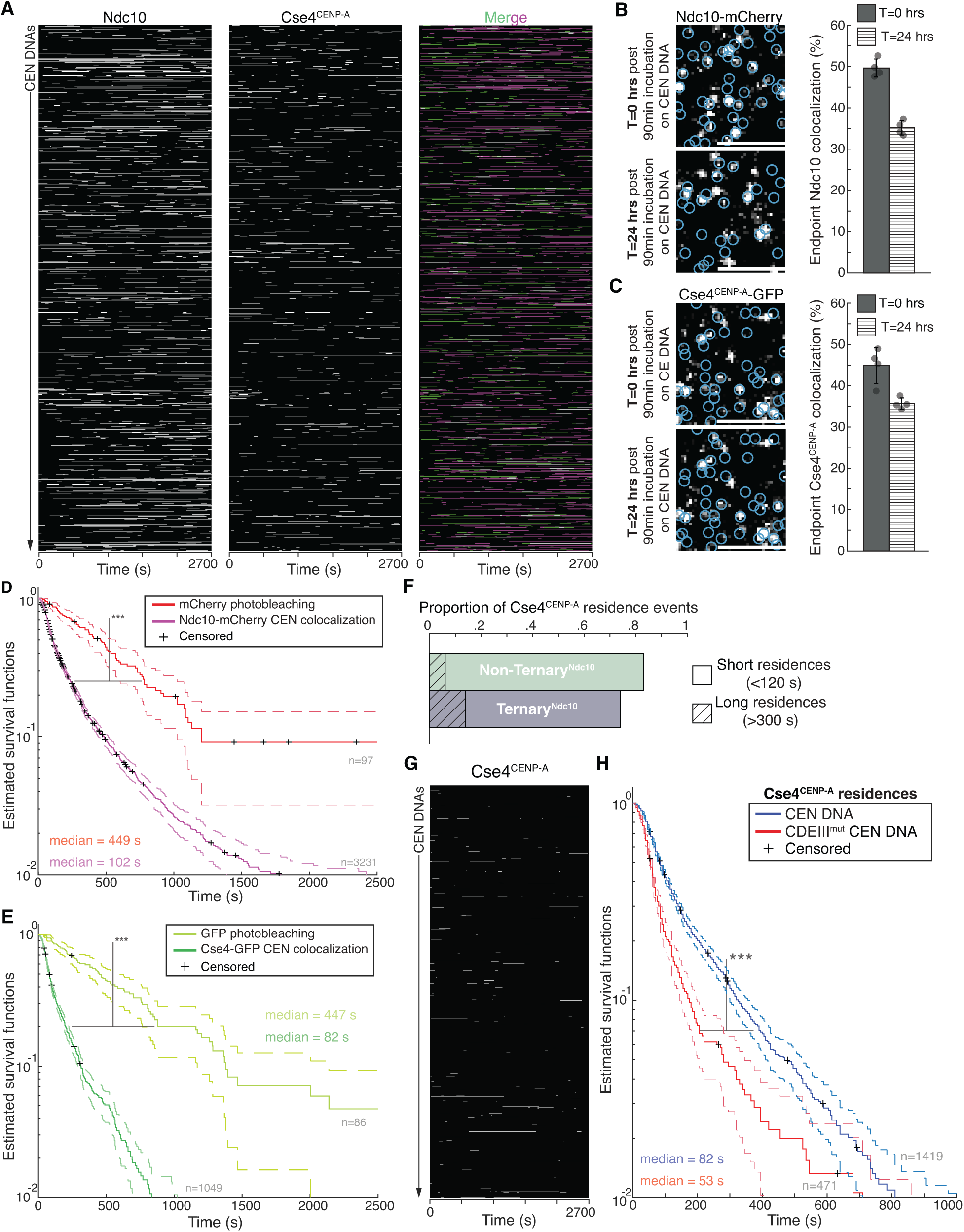
Ndc10 residence lifetimes are longer than those of Cse4^CENP-A^, which are not severely limited by photobleaching, while both are extremely stable on CEN DNA once removed from lysate. A Example plot of Ndc10 and Cse4^CENP-A^ residence pulses on CEN DNA identified via residence lifetime assays during an entire imaging sequence acquisition. Each row represents one identified CEN DNA with all identified residences shown over entire imaging sequence (2700 s) for Ndc10 (left) and Cse4^CENP-A^ (center) with merge of Ndc10 (magenta) and Cse4^CENP-A^ (green) indicating ternary residences (white). B Example images of TIRFM endpoint colocalization assays. Visualized Ndc10-mCherry on CEN DNA after 90 min incubation and removal of lysate (0 hrs -top panel) or after 24 hrs incubation at RT in imaging buffer (bottom panel) with colocalization shown in relation to identified CEN DNA in blue circles. Scale bars 3 μm. Graph shows quantification of Ndc10 endpoint colocalization on CEN DNA at 0 hrs and 24 hrs (50 ± 2.2%, 35 ± 1.7% respectively, avg ± s.d. n=4 experiments, each examining ∼1,000 DNA molecules from different extracts). C Example images of TIRFM endpoint colocalization assays. Visualized Cse4^CENP-A^ GFP on CEN DNA after 90 min incubation and removal of lysate (0 hrs -top panel) or after 24 hrs incubation at RT in imaging buffer (bottom panel) with colocalization shown in relation to identified CEN DNA in blue circles. Scale bars 3 μm. Graph shows quantification of Cse4^CENP-A^ endpoint colocalization on CEN DNA at 0 hrs and 24 hrs (45 ± 4.4%, 36 ± 1.4% respectively, avg ± s.d. n=4 experiments, each examining ∼1,000 DNA molecules from different extracts). D Kaplan-Meier analysis of mCherry photobleaching events (red median photobleaching lifetimes of 449 s (n=97)) and Ndc10 residences on CEN DNA (magenta median lifetime of 102 s (n=3231)). There was a significant difference between mCherry photobleaching lifetimes and Ndc10 residence lifetime survival plots (***) on CEN (two-tailed p-value of 0 as determined by log-rank test). E Kaplan-Meier analysis of GFP photobleaching events (yellow -median photobleaching lifetime of 447 s (n=86)) and Cse4^CENP-A^ residences on CEN DNA (red – median colocalization lifetime of 88 s (n=1054)). There was a significant difference between GFP photobleaching lifetimes and Cse4^CENP-A^ residence lifetime survival plots (***) on CEN DNA (two-tailed p-value of 0 as determined by log-rank test). 95% confidence intervals indicated (dashed lines), right-censored lifetimes (plus icons) were included and unweighted in survival function estimates. F Quantification of the proportion of short residences (<120 s) and long residences (>300 s) of **Non-Ternary^Ndc10^** Cse4^CENP-A^ residences (.77 and .06 respectively, n=612 over 3 experiments of ∼1000 DNA molecules using different extracts) or **Ternary^Ndc10^** Cse4^CENP-A^(.60 and .14 respectively, n=539 over 3 experiments of ∼1000 DNA molecules using different extracts). G Example plot of residences of Cse4^CENP-A^ on CDEIII^mut^ CEN DNA per imaging sequence. Each row represents one identified CEN DNA with all identified residences shown over entire imaging sequence (2700 s) for Cse4^CENP-A^. H Cse4^CENP-A^ residence lifetimes on CDEIII^mut^ CEN DNA are reduced. Estimated survival function plots of Kaplan-Meier analysis of residence lifetimes of Cse4^CENP-A^ on CEN DNA (blue – median lifetime of 82 s, n=1419 over 3 experiments of ∼1000 DNA molecules using different extracts) and residences on CDEIII^mut^ CEN DNA of Cse4^CENP-A^ (red of 52 s (n=471 over 3 experiments of ∼1000 DNA molecules using different extracts). There was a significant difference (***) between CEN DNA and CDEIII^mut^ CEN DNA lifetime survival plots (two-tailed p-value of 0 as determined by log-rank test). 95% confidence intervals indicated (dashed lines), right-censored lifetimes (plus icons) were included and unweighted in survival function estimates.

**Figure EV2.**
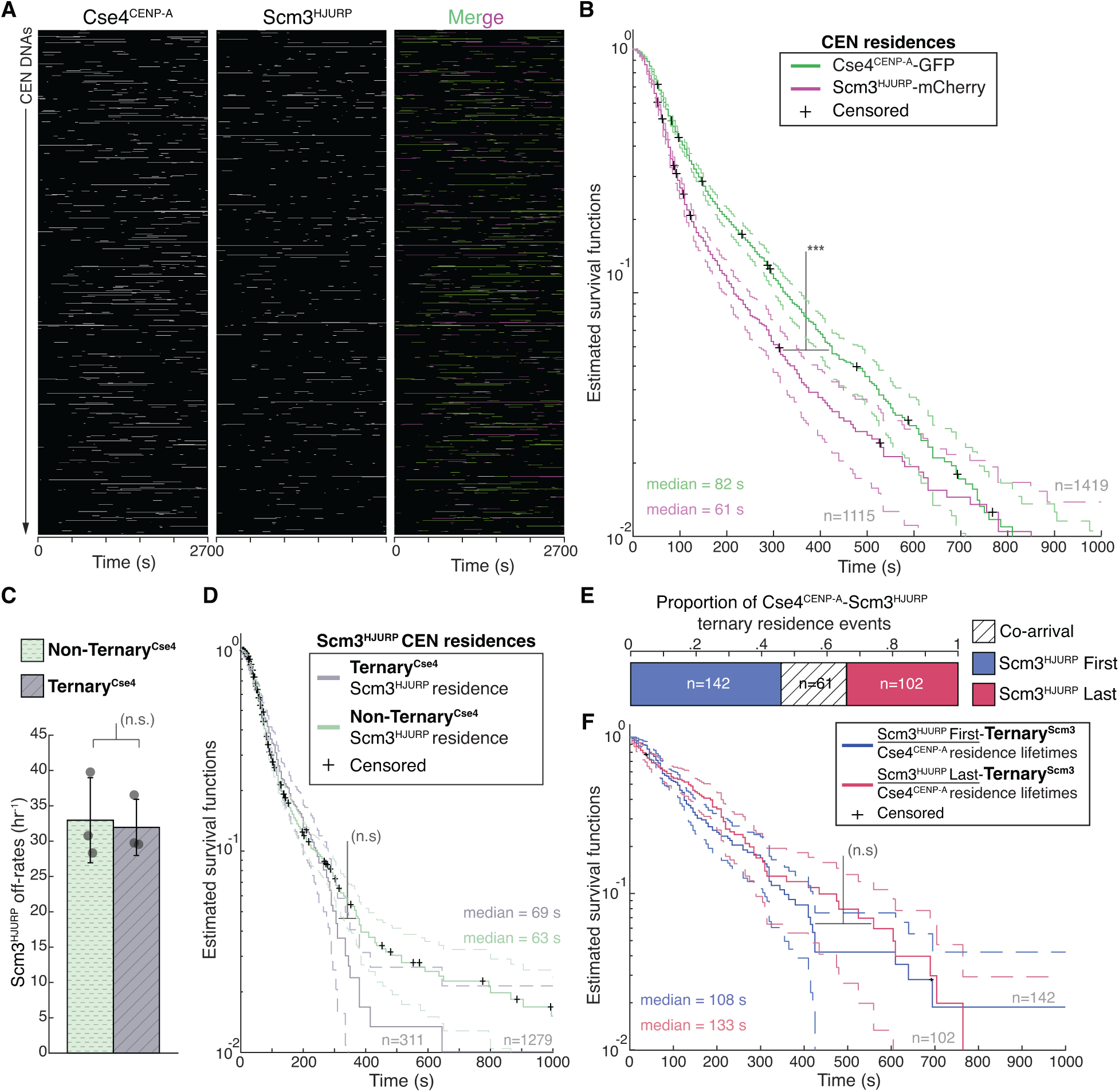
Scm3^HJURP^ has shorter residence lifetimes than Cse4^CENP-A^ on CEN DNA. A Example plot of Cse4^CENP-A^ and Scm3^HJURP^ residences on CEN DNA per imaging sequence. Each row represents one identified CEN DNA with all identified residences shown over entire imaging sequence (2700 s) Cse4^CENP-A^ (left) and Scm3^HJURP^ (center) with merge indicating Cse4^CENP-A^ (green), Scm3^HJURP^ (magenta) and ternary residences (white). B Estimated survival function plots of Kaplan-Meier analysis of Cse4^CENP-A^ residence lifetimes on CEN DNA (green – median lifetime of 82 s, n=1419 over 3 experiments of ∼1000 DNA molecules using different extracts), and residence lifetimes of Scm3^HJURP^ on CEN DNA (magenta – median lifetime of 61 s, n=1115 over 3 experiments of ∼1000 DNA molecules using different extracts). 95% confidence intervals indicated (dashed lines). Significant difference (***) between Cse4^CENP-A^ and Scm3^HJURP^ residence survival plots (two-tailed p-value of 9.1e-14 as determined by log-rank test), right-censored lifetimes (plus icons) were included and unweighted in survival function estimates. C Quantification of the estimated off-rates of Scm3^HJURP^ that never formed a ternary residence (**Non-Ternary^Cse4^**) and of Scm3^HJURP^ after ternary residence with Cse4^CENP-A^ (**Ternary^Cse4^**) on CEN DNA (114 s ± 13 s and 111 s ± 19 s respectively, avg ± s.d. n=2050 over 3 experiments of ∼1000 DNA molecules using different extracts). No significant difference between off-rates (n.s.) with a P-value of .87 as determined by two-tailed unpaired *t*-test. D. Estimated survival function plots of Kaplan-Meier analysis of the lifetimes of **Ternary^Cse4^** Scm3^HJURP^ residences on CEN DNA (purple – median lifetime of 69 s, n=311 over 3 experiments of ∼1000 DNA molecules using different extracts) and **Non Ternary^Cse4^** Scm3^HJURP^ residences on CEN DNA (green of 63 s, n=1279 over 3 experiments of ∼1000 DNA molecules using different extracts). No significant difference (n.s.) between **Ternary^Cse4^**and **Non-Ternary^Cse4^**survival plots (two-tailed p-value of .75 as determined by log-rank test). 95% confidence intervals indicated (dashed lines), right censored lifetimes (plus icons) were included and unweighted in survival function estimates. E Timing of all observed Cse4^CENP-A^ and Scm3^HJURP^ ternary residence events. The proportion when Scm3^HJURP^ precedes Cse4^CENP-A^ (<u>Scm3^HJURP^ First</u>) is 0.46, followed by.34 when Cse4^CENP-A^ precedes Scm3^HJURP^ (<u>Scm3^HJURP^ Last</u>), with a proportion of .20 co-arrival events (<u>Co-arrival</u> defined as residence initiation within 5 s of each, n=305 over 3 experiments of ∼1000 DNA molecules using different extracts). F Estimated survival function plots of Kaplan-Meier analysis of the lifetimes of <u>Scm3^HJURP^ First</u>-**Ternary^Scm3^** Cse4^CENP-A^ residences on CEN DNA (purple – median lifetime of 108 s, n=142 over 3 experiments of ∼1000 DNA molecules using different extracts) and <u>Scm3^HJURP^-Last</u>-**Ternary^Scm3^**Cse4^CENP-A^ residences on CEN DNA (green of 133 s, n=102 over 3 experiments of ∼1000 DNA molecules using different extracts). No significant difference (n.s.) between <u>Scm3^HJURP^-First and Scm3^HJURP^-Last</u> survival plots (two-tailed p-value of .59 as determined by log-rank test). 95% confidence intervals indicated (dashed lines), right-censored lifetimes (plus icons) were included and

**Figure EV3.**
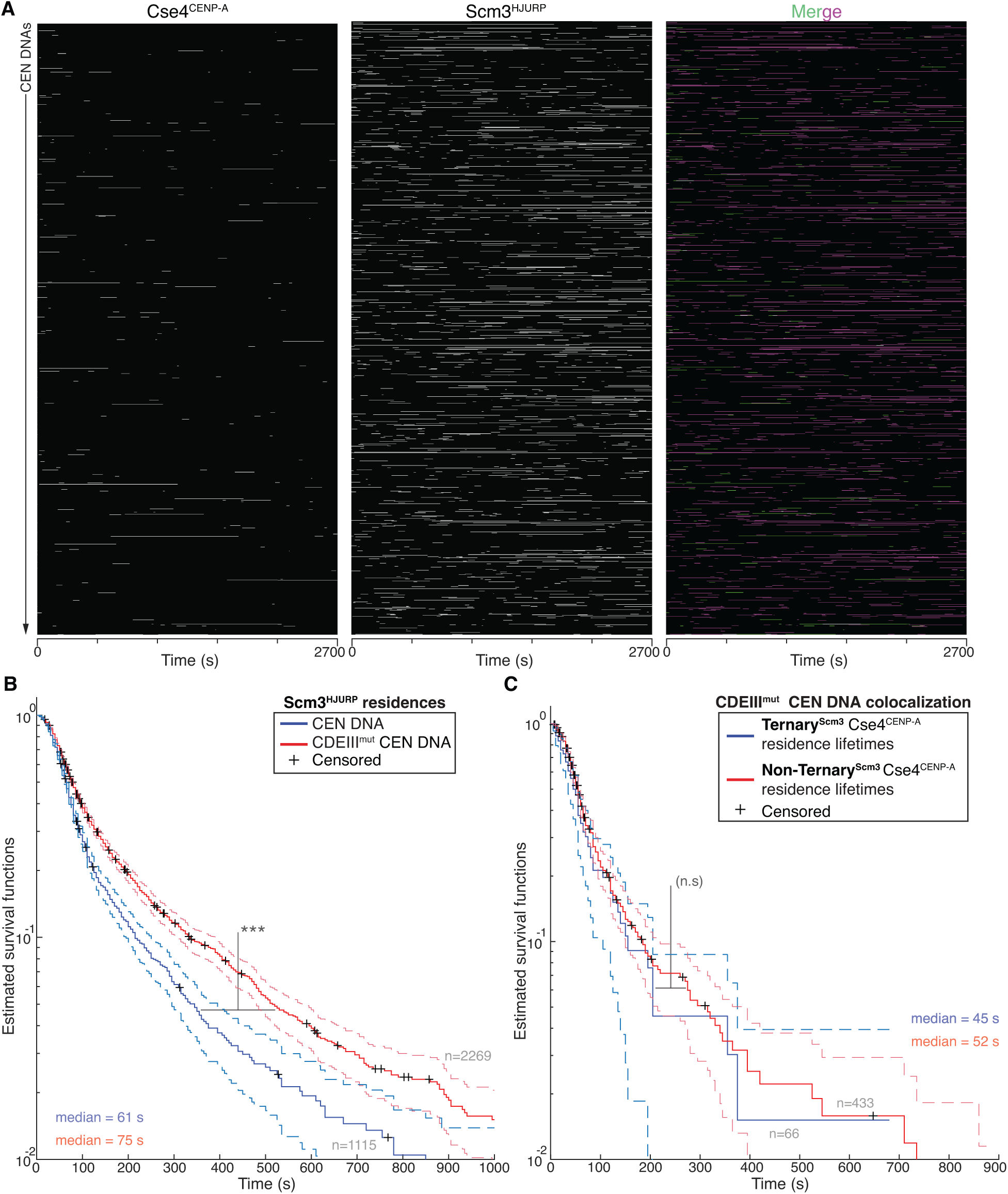
Cse4^CENP-A^ interacts transiently with CDEIII^mut^ CEN DNA with or without its chaperone Scm3^HJURP^. A Example plot of residences of Cse4^CENP-A^ and Scm3^HJURP^ on CDEIII^mut^ CEN DNA per imaging sequence. Each row represents one identified CEN DNA with all identified residences shown over entire imaging sequence (2700 s) for Cse4^CENP-A^ (left) and Scm3^HJURP^ (center) with merge indicating Cse4^CENP-A^ (green), Scm3^HJURP^ (magenta) and ternary residences (white). B Scm3^HJURP^ residences are longer on CDEIII^mut^ CEN DNA. Estimated survival function plots of Kaplan-Meier analysis of Scm3^HJURP^ residence lifetimes on CEN DNA (blue median lifetime of 61 s, n=1115 over 3 experiments of ∼1000 DNA molecules using different extracts), and residence lifetimes of Scm3^HJURP^ on CDEIII^mut^ CEN DNA (red median lifetime of 75 s, n=2269 over 3 experiments of ∼1000 DNA molecules using different extracts). There was a significant difference (***) between CEN DNA and CDEIII^mut^ CEN DNA lifetime survival plots (two-tailed p-value of 5.13e-13 as determined by log-rank test). 95% confidence intervals indicated (dashed lines), right-censored lifetimes (plus icons) were included and unweighted in survival function estimates. C Cse4^CENP-A^ residence lifetimes are similar without Scm3^HJURP^ a on CDEIII^mut^ CEN DNA. Estimated survival function plots of Kaplan-Meier analysis of ternary residence lifetimes of **Ternary^Scm3^** Cse4^CENP-A^ residences on CDEIII^mut^ CEN DNA (blue – median lifetime of 45 s, n=66 over 3 experiments of ∼1000 DNA molecules using different extracts) and **Non-Ternary^Scm3^** Cse4^CENP-A^ residences on CDEIII^mut^ CEN DNA (red of 52 s (n=433 over 3 experiments of ∼1000 DNA molecules using different extracts). There was no significant difference (n.s.) between **Non-Ternary^Scm3^** and **Ternary^Scm3^** survival plots (two-tailed p-value of .27 as determined by log-rank test). 95% confidence intervals indicated (dashed lines), right-censored lifetimes (plus icons) were included and unweighted in survival function estimates.

**Figure EV4.**
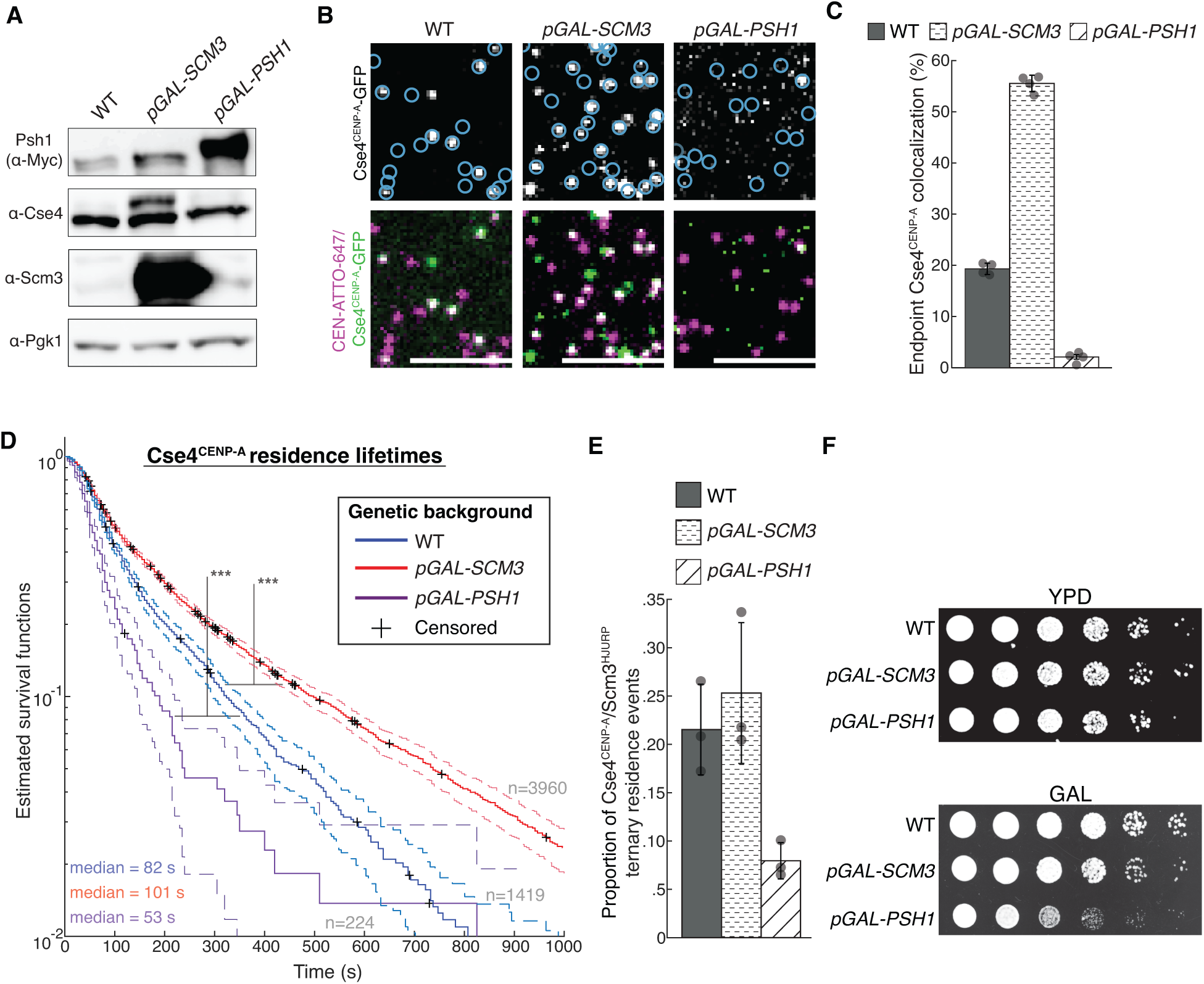
Scm3^HJURP^-Cse4^CENP-A^ complex is limiting for stable centromeric association of Cse4^CENP-A^. A Immunoblot analysis of whole cell extracts from WT, *pGAL-SCM3* and *pGAL-PSH1* cells using indicated antibodies (all panels cropped from the same blot). B Example images of TIRFM endpoint colocalization assays. Top panels show visualized Cse4^CENP-A^ -GFP on CEN DNA in extracts from a WT genetic background (top-left panel) or extracts containing overexpressed Scm3^HJURP^ (*pGAL-SCM3,* top-middle panel) or overexpressed Psh1 (*pGAL-PSH1,* top-right panel) with colocalization shown in relation to identified CEN DNAs in blue circles. Bottom panels show overlay of DNA channel (magenta) with Cse4^CENP-A^ -GFP (green). C Quantification of endpoint colocalization of Cse4^CENP-A^ on CEN DNA in extracts from a WT genetic background, extracts that contain overexpressed Scm3^HJURP^ or extracts that contain overexpressed Psh1 (19 ± 1.1%, 56 ± 1.6% and 2.1 ± 0.4% respectively, avg ± s.d. n=4 experiments, each examining ∼1,000 DNA molecules from different extracts). Scale bars 3 μm. D Estimated survival function plots of Kaplan-Meier analysis of residence lifetimes of Cse4^CENP-A^ on CEN DNA in extracts from WT genetic background (blue median lifetime of 82 s, n=1419 over 3 experiments of ∼1000 DNA molecules using different extracts), or from extracts that contain overexpressed Scm3^HJURP^ (red – median lifetime of 101 sec, n=3960 over 3 experiments of ∼1000 DNA molecules using different extracts) or extracts that contain overexpressed Psh1 (purple – median lifetime of 52 sec, n=224 over 3 experiments of ∼1000 DNA molecules using different extracts). Significant difference (***) between survival plots in WT extracts compared to those overexpressing Psh1 (two-tailed p-value of 8.9e-11 as determined by log-rank test) or Scm3^HJURP^ (two-tailed p-value of 0 as determined by log-rank test). 95% confidence intervals indicated (dashed lines), right censored lifetimes (plus icons) were included and unweighted in survival function estimates. E Proportion of ternary residences of Cse4^CENP-A^ with Scm3^HJURP^ on CEN DNA for WT extracts (0.22 ± .05, avg ± s.d. n=1419 over 3 experiments of ∼1000 DNA molecules using different extracts), extracts containing overexpressed Scm3^HJURP^ (0.25 ± .07, avg ± s.d. n=3960 over 3 experiments of ∼1000 DNA molecules using different extracts) and extracts containing overexpressed Psh1 (0.08 ± .02, avg ± s.d. n=224 over 3 experiments of ∼1000 DNA molecules using different extracts). (**F**) Serial 5-fold dilutions of the following yeast strains were plated and grown two days on YPD and three days on galactose (GAL) at 23° C: WT (SBY21441), *pGAL-SCM3* (SBY21443), and *pGAL-PSH1* (SBY20836).

**Figure EV5.**
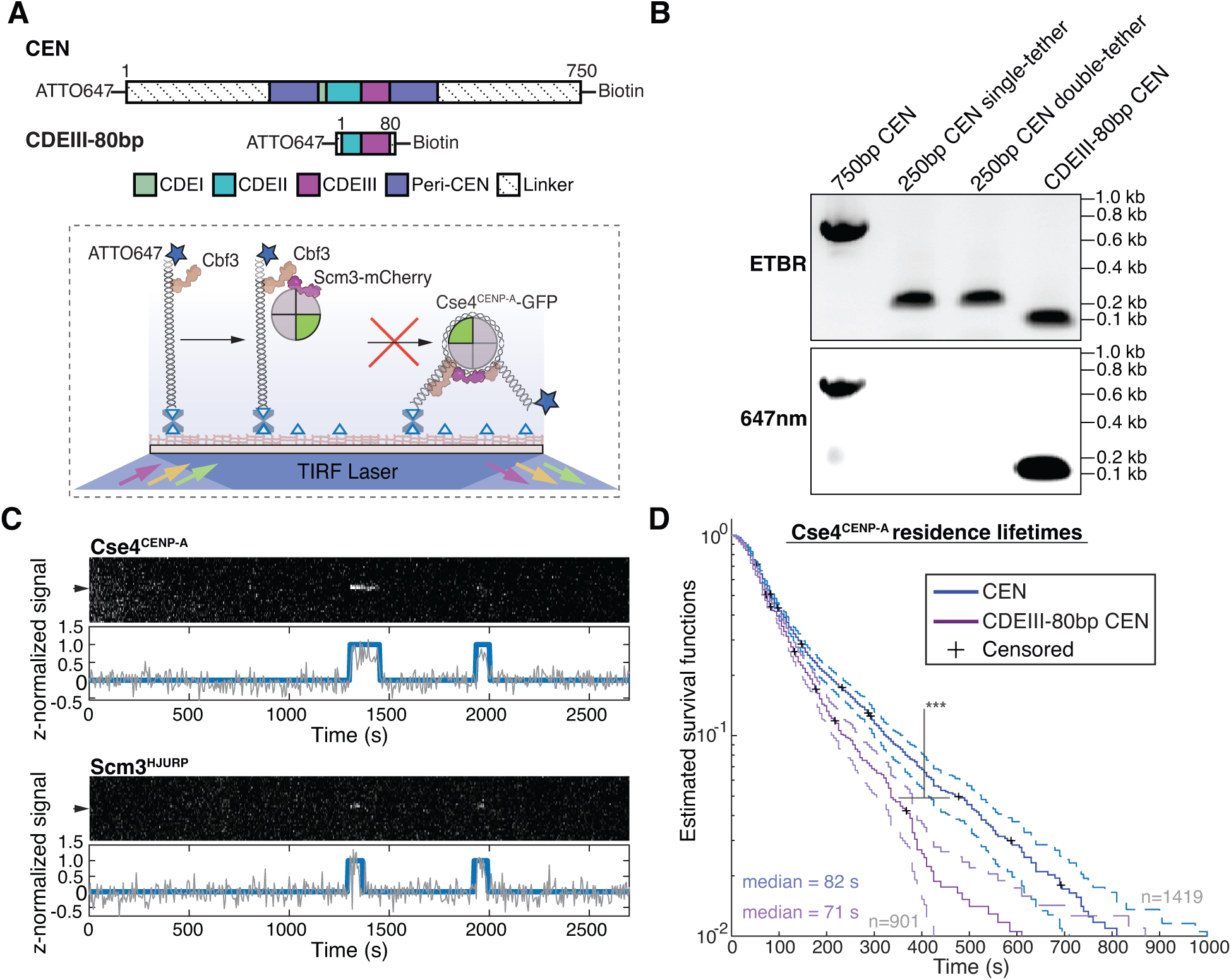
Cse4^CENP-A^ residence lifetimes are significantly reduced on CDEIII-80 bp mutant CEN DNA. A Schematic of overview of CDEIII-80 bp mutant CEN DNA, the canonical CEN DNA is shortened to 80 bp to prevent nucleosome formation and then similarly functionalized to the coverslip via a single biotin at the 5’end and functionalized with an organic dye at the free 3’ end. B CEN assembly templates including WT 750 bp CEN DNA, 250 bp single-tether CEN DNA, 250 bp double-tethered CEN DNA and CDEIII-80 bp CEN DNA as visualized via EtBr (top panel) or 647nm excitation (bottom panel) on a 1% agarose gel. C Representative colocalization traces of Cse4^CENP-A^ and Scm3^HJURP^ on a single CDEIII 80 bp CEN DNA. Top panel includes kymograph of Cse4^CENP-A^ (top-488 nm) in relation to single identified CEN DNA (arrow), with normalized intensity trace (grey-bottom) as well as identified colocalization pulses (blue). Bottom panel includes kymograph of Scm3^HJURP^ (bottom-568 nm) in relation to the same identified CEN DNA (arrow), with normalized intensity trace (grey-bottom) as well as identified colocalization pulse (blue). Cases where identified pulses in Scm3^HJURP^ and Cse4^CENP-A^ coincide represent observed colocalization of both proteins on single CDEIII-80 bp CEN DNA. Images acquired every 5 seconds with normalized fluorescence intensity shown in arbitrary units. D Estimated survival function plots of Kaplan-Meier analysis of all identified CEN DNA colocalization events of Cse4^CENP-A^ (blue median lifetime of 82 s, n=1619 over 3 experiments of ∼1000 DNA molecules using different extracts) and identified colocalization events on CDEIII-80 bp CEN DNA of Cse4^CENP-A^ (red – median lifetime of 71 s, n=901 over 3 experiments of ∼1000 DNA molecules using different extracts). Significant difference (***) between CEN DNA and CDEIII-80 bp CEN DNA lifetime survival plots (two-tailed p-value of 1.0e-6 as determined by log-rank test). 95% confidence intervals indicated (dashed lines), right-censored lifetimes (plus icons) were sincluded and unweighted in survival function estimates.

## Notes

https://github.com/FredHutch/Automated-Single-Molecule-Colocalization-Analysis

### Competing Interest Statement

The authors have declared no competing interest.

### Summary of Updates

This is a revised version of the manuscript that addresses the reviewers' comments as described in the rebuttal.

