## Appendix for "Direct observation of coordinated assembly of individual native centromeric nucleosomes"

**Table of Contents**

**Appendix Supplemental Methods……………………………………..………………………..3**

**Appendix Supplemental Methods References……………………..………………………..3**

**Expanded View Video 1:** Ndc10 association with CEN3 DNA preceding Cse4$^{CENP-A}$ recruitment…......………………………………………………………………………..………..4

**Expanded View Video 2:** Ndc10 association with CEN3 DNA after Cse4$^{CENP-A}$ recruitment…......………………………………………………………………………...……….4

**Expanded View Video 3:** Stable Cse4$^{CENP-A}$ recruitment coincides with ternary CEN3 DNA residence with Scm3$^{HJURP}$……………………………..…………………………………...…4

**Expanded View Video 3:** Double-tethered 250 bp CEN3 DNA fails to stably recruit Cse4$^{CENP-A}$…………………………………………………...…...…………………..4

**Expanded View Video 4:** 80 bp CDEIII mutant CEN DNA fails to stably recruit Cse4$^{CENP-A}$….4

**Appendix Figure S1.** Two copies of Ndc10 and Cse4$^{CENP-A}$ associate with CEN3 in TIRFM endpoint colocalization assays..…………………………...……………………………...…….5

**Appendix Figure S2.** Cse4$^{CENP-A}$ behaves similarly on CEN template DNA from different chromosomes**…..…………………………...…………...………...……………………………….6**

**Appendix Figure S3.** Post Scm3$^{HJURP}$-Cse4$^{CENP-A}$ estimated off-rates are significantly reduced on "very unstable" CDEII mutant CEN DNA**…...…………………………………..……………….8**

**Appendix Figure S4.** Levels of H3 incorporation on are low on CEN3 or 601-hybrid CEN template DNAs but increase in the absence of the CBF3 complex on the Widom 601 targeting sequence**.………...…………………………………………..………………………….....9**

**Appendix Table 1:** Related to Figures 1-6. List of *S. cerevisiae* strains used in this study**………..........................................……………………………………………...…….11**

**Appendix Table 2:** Related to Figures 1-6. Plasmids used to generate *S. cerevisiae* strains and for CEN DNA template generation**………..................................……………………………………………...….12**

**Appendix Table 3:** Related to Figures 1-6. DNA oligonucleotides used in this study for *S. cerevisiae* strain construction and CEN DNA template sequence generation**…………………...13**

**Appendix Supplemental Methods:**

**Photobleaching assays**

For the single molecule photobleaching assays, surface bound CEN3 DNA-containing flow chambers were filled with 100 μL of whole cell extract (WCE) containing protein of interest via pipetting and wicking with filter paper. After addition of WCE, slides were incubated for 90 min at 25°C and then WCE was washed away with Buffer L. Flow chambers were then filled with Buffer L for imaging. Time-lapse images were started with single acquisition of CEN3 DNA (647 nm exposure) followed by continuous acquisition with 200-ms exposure until the field of view was bleached in the channel of interest (488 nm for $Cse4^{CENP\text{-}A}$-GFP and 568 nm for Ndc10-mCherry). CEN3 DNA-associated proteins were then identified and the number of photobleaching steps of individual CEN3-bound $Cse4^{CENP\text{-}A}$-GFP or Ndc10-mCherry was obtained by tracking the fluorescence intensity in FIJI(Schindelin *et al*, 2012).

For the single molecule lifetime photobleaching approximations, WCE containing either $Cse4^{CENP\text{-}A}$-GFP or Ndc10-mCherry was introduced to a non-passivated flow chamber and incubated for 15 min at 25°C to permit non-specific binding to coverslip surface. After incubation WCE was washed away with Buffer L followed by introduction of 100 μL of WCE containing no fluorescently tagged proteins (SBY3). After introduction of WCE, acquisition was started immediately for $Cse4^{CENP\text{-}A}$-GFP at 488 nm for 200 ms exposures every 5 s, and for Ndc10-mCherry at 561 nm for 200 ms exposures every 5 s. Acquisition was performed until the field of view was bleached in channel of interest. The fluorescence lifetimes of individual $Cse4^{CENP\text{-}A}$-GFP or Ndc10-mCherry were tracked in FIJI followed by Kaplan-Meier analysis performed in MATLAB (R2021a).

**Appendix Supplemental Videos:**

**Appendix Video S1. Ndc10 association with CEN3 DNA preceding stable Cse4$^{CENP\text{-}A}$ recruitment.** Movie showing the colocalization to single CEN3 DNA (647nm, center of left panel) of Cse4$^{CENP\text{-}A}$-GFP (488 nm, middle panel) and Ndc10-mCherry (568 nm, right panel). This movie corresponds to Figure 2B. Scale bar 3 μm.

**Appendix Video S2. Ndc10 association with CEN3 DNA after stable Cse4$^{CENP\text{-}A}$ recruitment.** Movie showing the colocalization to single CEN3 DNA (647nm, center of left panel) of Cse4$^{CENP\text{-}A}$-GFP (488 nm, midle panel) and Ndc10-mCherry (568 nm, right panel). This movie corresponds to Figure 2B. Scale bar 3 μm.

**Appendix Video S3: Stable Cse4$^{CENP\text{-}A}$ recruitment coincides with ternary CEN3 DNA residence with Scm3$^{HJURP}$**. Movie showing the colocalization to single CEN3 DNA (647nm, left panel) of Cse4$^{CENPA}$-GFP (488 nm, middle panel) and Scm3$^{HJURP}$-mCherry (568 nm, right panel). This movie corresponds to Figure 3A. Scale bar 3 μm.

**Appendix Video S4: Double-tethered 250 bp CEN3 DNA fails to stably recruit Cse4$^{CENP\text{-}A}$.** Movie showing the colocalization to single Ndc10-mCherry on single tethered CEN3 DNA (568 nm, top left panel) of Cse4$^{CENPA}$-GFP (488 nm, top right panel) or on double tethered CEN3 DNA (568 nm, center of bottom left panel) of Cse4$^{CENPA}$-GFP (488 nm, bottom right panel). This movie corresponds to Figure 4E (top and middle panels). Scale bar 3 μm.

**Appendix Video S5: 80 bp CDEIII mutant CEN DNA fails to stably recruit Cse4$^{CENP\text{-}A}$.** Movie showing the colocalization to single 80 bp CDEIII mutant CEN DNA (647nm, center of left panel) of Cse4$^{CENP\text{-}A}$-GFP (488 nm, right panel). This movie corresponds to Figure 4E (bottom panel). Scale bar 3 μm.

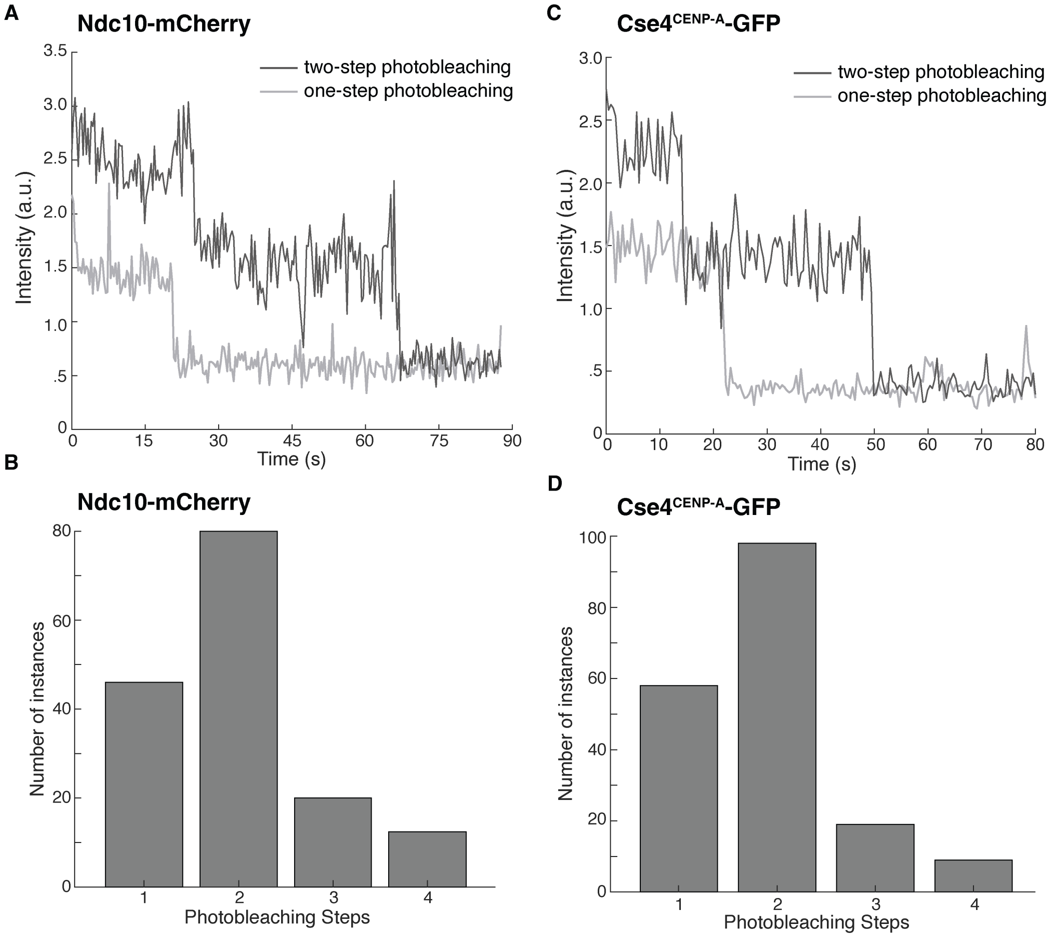


**Appendix Figure S1: Two copies of Ndc10 and Cse4$^{CENP\text{-}A}$ associate with CEN3 in TIRFM endpoint colocalization assays.** (**A**) Representative photobleaching traces of Ndc10-mCherry fluorescence on colocalized CEN3 DNA. (**B**) Histogram of fluorescence photobleaching steps of Ndc10-mCherry (n = 155). (**C**) Representative photobleaching traces of Cse4$^{CENP\text{-}A}$-GFP fluorescence on colocalized CEN3 DNA. (**D**) Histogram of fluorescence photobleaching steps of Cse4$^{CENP\text{-}A}$-GFP (n = 184).

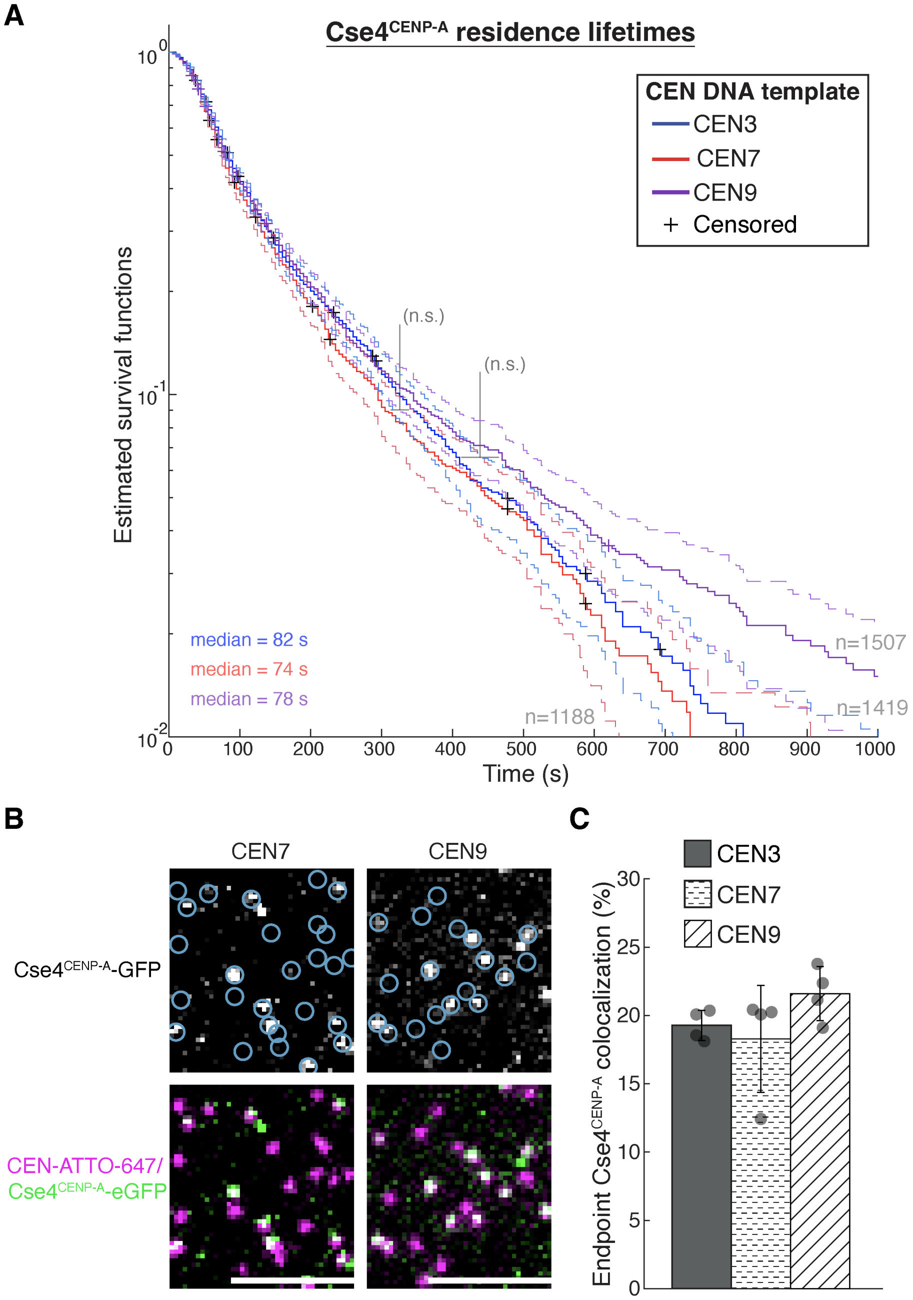


**Appendix Figure S2: Cse4$^{CENP\text{-}A}$ behaves similarly on CEN template DNA from different chromosomes.** (**A**) Estimated survival function plots of Kaplan-Meier analysis of Cse4$^{CENP\text{-}A}$ residence lifetimes on CEN3 DNA (blue - median lifetime of 82 sec, n=1614 over 3 experiments of ~1000 DNA molecules using different extracts), CEN7 DNA (red - median lifetime of 74 sec, n=1473 over 3 experiments of ~1000 DNA molecules using different extracts) and CEN9 DNA (purple - median lifetime of 78 s, n=2242 over 3 experiments of ~1000 DNA molecules using different extracts). No significant difference (n.s) between Cse4$^{CENP\text{-}A}$ on CEN3 DNA and CEN9

DNA residence lifetime survival plots (two-tailed p-value of .62 as determined by log-rank test) or compared to CEN7 DNA (two-tailed p-value of .053 as determined by log-rank test). 95% confidence intervals indicated (dashed lines), right-censored lifetimes (plus icons) were included and unweighted in survival function estimates. (**B**) Example images of TIRFM endpoint colocalization assays. Top panels show visualized $Cse4^{CENP-A}$ -GFP in lysates containing $Scm3^{HJURP}$-mCherry on CEN7 DNA template (top-left panel) or CEN9 DNA template (top-right panel) with colocalization shown in relation to identified CEN DNAs in blue circles. Bottom panels show overlay of DNA channel (magenta) with $Cse4^{CENP-A}$-GFP (green) on CEN7 DNA (bottom-left panel) or CEN9 DNA (bottom-right panel. (**C**) Quantification of endpoint colocalization of $Cse4^{CENP-A}$ on CEN3 DNA, CEN7 DNA and on CEN9 (19 ± 1.1%, 18 ± 3.9% and 22 ± 2.0% respectively, avg ± s.d. n=4 experiments, each examining ~1,000 DNA molecules from different extracts). Scale bars 3 μM.

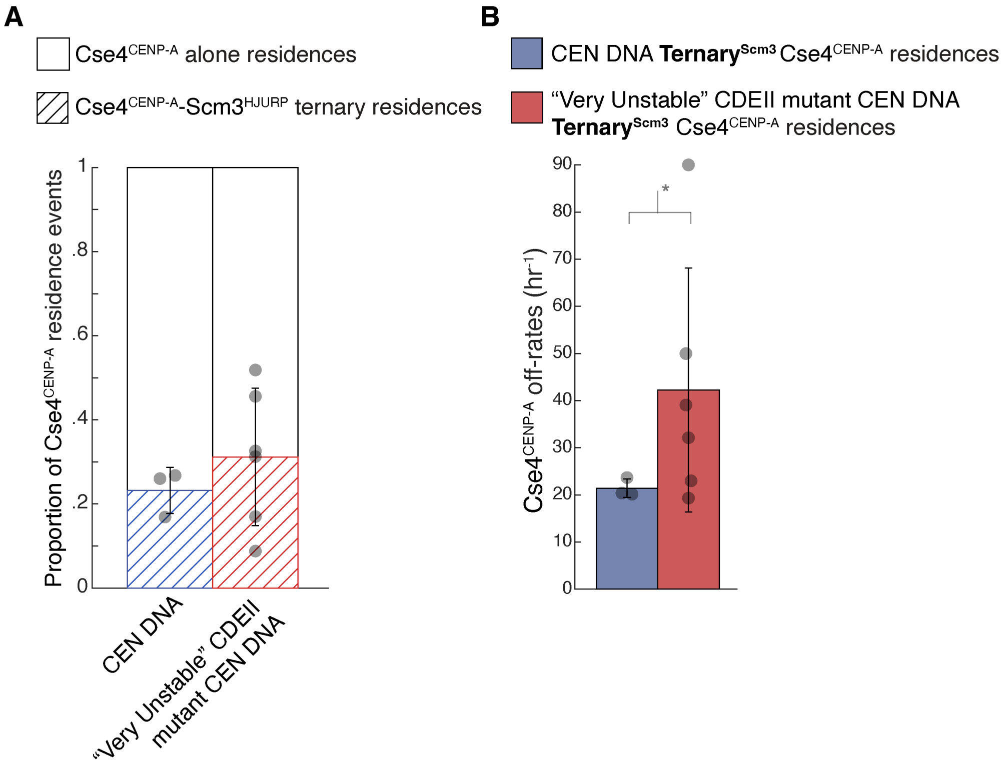


**Appendix Figure S3: Ternary$^{Scm3}$ Cse4$^{CENP-A}$ residence off-rates are significantly increased on "very unstable" CDEII mutant CEN DNA**. (**A**) Proportion of ternary residences of Cse4$^{CENP-A}$ with Scm3$^{HJURP}$ on CEN3 DNA (blue - 0.23 ± .06, avg ± s.d. n=1863 over 3 experiments of ~1000 DNA molecules using different extracts), or on "very unstable" CDEII mutant CEN DNA (red - 0.31 ± .16, avg ± s.d. n=2562 over 3 experiments of ~1000 DNA molecules using different extracts). (**B**) Cse4$^{CENP-A}$ has slower off-rates after colocalization with Scm3$^{HJURP}$ on CEN DNA than on "Very Unstable" CDEII mutant CEN DNA. Quantification of the estimated off-rates of **Ternary$^{Scm3}$** Cse4$^{CENP-A}$ residences on CEN DNA (21 hr$^{-1}$ ± 2 hr$^{-1}$, avg ± s.d. n=305 over 3 experiments of ~1000 DNA molecules using different extracts) and **Ternary$^{Scm3}$** Cse4$^{CENP-A}$ residences on "Very Unstable" CDEII mutant CEN DNA (42 hr$^{-1}$ ± 26 hr$^{-1}$, avg ± s.d. n=358 over 6 experiments of ~1000 DNA molecules using different extracts). Significant difference between off-rates (*) with a P-value of .046 as determined by two-tailed unpaired *t*-test.

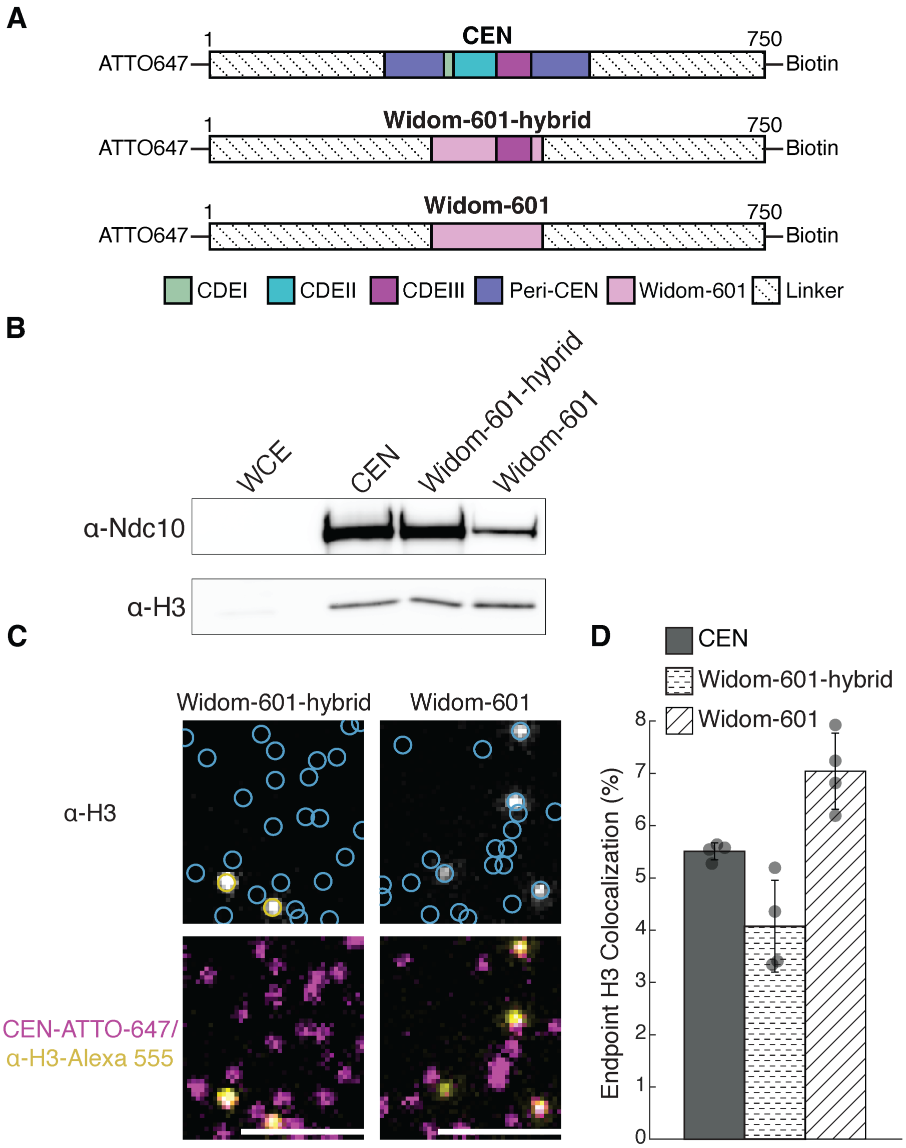


**Appendix Figure S4: Levels of H3 incorporation on are low on CEN3 or Widom 601-hybrid CEN template DNAs but increase in the absence of the CBF3 complex on the Widom 601 targeting sequence.** (**A**) Schematic of centromeric templates. (**B**) Immunoblot analysis of kinetochore assembly on WT, Widom-601-hybrid, and Widom-601 templates from WT whole cell extracts (WCE). Immunoblotted with indicated antibodies. Ndc10 could be visualized in WCE at much longer exposures (not shown). (**C**) Example images of TIRFM endpoint colocalization imaging. Top panels show immunofluorescence of H3 on Widom-601-hybrid CEN3 DNA (top-left panel) or Widom-601 DNA (top-right panel) with colocalization shown in relation to identified CEN3 DNAs in blue circles. Bottom panels show overlay of DNA channel (magenta) with H3 (yellow). (**D**) Quantification of endpoint colocalization of H3 on WT CEN3 DNA, Widom-601-hybrid CEN DNA and Widom-601 DNA (5.5 ± 0.2%, 4.1 ± 0.9% and 7.1 ± 0.7% respectively, avg ± s.d. avg ± s.d. n=4 experiments, each examining ~1,000 DNA molecules from different extracts). Scale bars 3 μm.

**Appendix Table 1: Related to Figures 1-6. List of *S. cerevisiae* strains used in this study.**

| Strain | Genotype |
|---|---|
| | All strains derived from W303 |
| SBY3 | *MAT***a** *ura3-1 leu2,3-112 his3-11 trp1-1 ade2-1 LYS2 can1-100 bar1-1* |
| SBY8315 | *MAT***a** *ura3-1 leu2,3-112 his3-11 trp1-1 ade2-1 can1-100 bar1 NDC10-mCherry:HPH* |
| SBY19926 | *MAT***a** *ura3-1::pCSE4-CSE4-XbaI(GFP):URA3 leu2,3-112 his3-11 trp1-1 ade2-1 LYS2+ can1-100 bar1 cse4Δ::KanMX* |
| SBY20464 | *MATα ura3-1::pCSE4-CSE4-XbaI(GFP):URA3 leu2,3-112 his3-11 trp1-1 ade2-1 LYS2 can1-100 bar1 cse4Δ::KanMX SCM3-mCherry:HPH* |
| SBY20629 | *MATα ura3-1::pCSE4-CSE4-XbaI(GFP):URA3 leu2,3-112 his3-11::pGPD1-OsTIR1:HIS3 trp1-1 ade2-1 LYS2+ can1-100 bar1 cse4Δ::KanMX SCM3-mCherry:HPH okp1-3V5-IAA7:KanMX6* |
| SBY20836 | *MATα ura3-1::pCSE4-CSE4-XbaI(GFP):URA3 leu2,3-112::pGAL-13MYC-PSH1:LEU2 his3-11 trp1-1 ade2-1 LYS2 can1-100 bar1 cse4Δ::KanMX SCM3-mCherry:HPH* |
| SBY21110 | *MATα ura3-1::pCSE4-CSE4-XbaI(GFP):URA3 leu2,3-112 his3-11 trp1-1 ade2-1 LYS2 can1-100 bar1 cse4Δ::KanMX NDC10-mCherry:HPH* |
| SBY21361 | *MATα ura3-1::pCSE4-CSE4-XbaI(GFP):URA3 leu2,3-112 his3-11 trp1-1 ade2-1 LYS2+ can1-100 bar1 cse4Δ::KanMX SCM3-mCherry:HPH chl4-K13S-13Myc:HIS3* |
| SBY21441 | *MATα ura3-1::pCSE4-CSE4-XbaI(GFP):URA3 leu2,3-112 his3-11 trp1-1 ade2-1 LYS2 can1-100 bar1 cse4Δ::KanMX SCM3-mCherry:HPH PSH1-13MYC:HIS* |
| SBY21443 | *MATα ura3-1::pCSE4-CSE4-XbaI(GFP):URA3 leu2,3-112::pGAL-SCM3-mCherry:LEU2 his3-11 trp1-1 ade2-1 LYS2 can1-100 bar1 cse4Δ::KanMX SCM3-mCherry:HPH PSH1-13MYC:HIS* |

**Appendix Table 2: Related to Figures 1-6. Plasmids used to generate *S. cerevisiae* strains and for CEN DNA template generation.**

| Plasmid | Description | Source |
|---|---|---|
| pSB64 | *13MYC, HIS3MX6* | Biggins Lab |
| pSB963 | *WT CEN3, 8LacO, TRP1* | Biggins Lab |
| pSB972 | *Mutant CEN3 (CCG->AGC CEN mutant), 8 LacO, TRP1* | Biggins Lab |
| pSB1582 | *mCherry, HPH* | Biggins Lab |
| pSB1617 | *pCSE4-CSE4(1-80)-GFP-(81-229), URA3* (integrating) | Biggins Lab |
| pSB2066 | *3V5-IAA7, KanMX6* | Biggins Lab |
| pSB2273 | *pGPD1-OsTIR1, HIS3* (integrating) | Biggins Lab |
| pSB2887 | *Widom-601* | Biggins Lab |
| pSB2953 | *CEN7, HIS3* | Biggins Lab |
| pSB3167 | *pSCM3-SCM3-mCherry HPH, LEU (integrating)* | Biggins Lab |
| pSB3252 | *pGAL-SCM3-mCherry*, *LEU2* (integrating) | Biggins Lab |
| pSB3264 | *Widom-601-CDEIII (hybrid)* | Biggins Lab |
| pSB3288 | *pGAL-13MYC-PSH1, LEU2* (integrating) | Biggins Lab |
| pSB3335 | *CEN3-CDEII-α-sat, 8LacO, TRP1* | Biggins Lab |
| pSB3336 | *CEN3-CDEII-H1, 8LacO, TRP1* | Biggins Lab |
| pSB3337 | *CEN3-CDEII-L1, 8LacO, TRP1* | Biggins Lab |
| pSB3338 | *CEN9, 8LacO, TRP1* | Biggins Lab |
| pSB3415 | *CEN3-CDEII-L5, 8LacO, TRP1* | Biggins Lab |
| pSB3416 | *CEN3-CDEII-H14, 8LacO, TRP1* | Biggins Lab |

**Appendix Table 3: Related to Figures 1-6. DNA oligonucleotides used in this study for *S. cerevisiae* strain construction and CEN DNA template sequence generation.**

| Primer | Sequence | Purpose |
|---|---|---|
| SB51 | 5' - GGAGGCATGACCATCAAAATTCATTTGATGGTCTGTTAGTATATCTATCTAAC CGGATCC CCGGGT TAATTAA- 3' | 5' primer to tag *NDC10* |
| SB52 | 5' - CATACATGTCGGTATCCCTATACGAAACAGTTTAAACTTCGAAGCTCCCTCA GA ATTCGAGCTCGT TTAAAC- 3' | 3' primer to tag *NDC10* |
| SB64 | 5' - CCGAAAAAGGGAAAAATCGGCTCCAGCCCTGAAGCACAAATATCACTATCGA TGAATTCGAGCTCGTT- 3' | 3' primer to delete *CSE4* |
| SB67 | 5' - CAGAAGAAGGACTGAATATAGAAAGAATACTAATATAACATAATCCGGATCC CCGGGTTAATTAA- 3' | 5' primer to delete *CSE4* |
| SB1713 | 5' - TTGTCCACTTGTTGCCACTGA- 3' | 3' primer to tag *SCM3* with mCherry with pSB3167 |
| SB1871 | 5' - CAACCGAAAAGGCAGAAACGCTTCCGTGTAGTTCTAGGAGACAGTGACGAT GAAGGTCGACGGATCCCCGGGTT- 3' | 5' primer to tag *PSH1* |
| SB1872 | 5' - CATAAAAGTTCCGTACATATGCCGTTCCCGCTAGTTAAAATGTTCAGATTACG ATGAATTCGAGCTCGTT- 3' | 3' primer to tag *PSH1* |
| SB4549 | 5' - GCACCATGAGTCGCACCAAGATAAGACCGAAGAAGATATACACCGGATCCC CGGGTTAATTAA- 3' | 5' primer to tag *OKP1* |

| | | |
|---|---|---|
| SB4550 | 5' - CAAATATTTAGTTATATGCATCGTAATCGTAAACTCTGAAACAATGGATTATC GGAATTCGAGCTCGTTTAAAC- 3' | 3' primer to tag *OKP1* |
| SB6572 | 5' - TATGTACCAACTTCCGACACACTAGTAGTTTTCtcACAATTGATGAgtCTGCCG GTAACGGTATTATATGATCTTACGCTATCATGGTTCGCAAAATTCGGTGGGT CATTTGATGGTGACATATATTTATTGACAGAAACATTAGACTTACTGATTGAG AAAGGCGTGAGtCGAAATGTTATAGTAAATAGGATATTATACGTATACTGGCC GGATGGCCTGAATGTTTTCCAATTAGCAGAAATAGATTGCCATTTAATGATAA GTtcACCAGAGtcATTTAgtTGGCTTCCATCAAAAGCTTTACGGGGAGATGGGA AACCTTACGTAGTAAAGCTCCAACCTGCCAAATTTATAGAAAATTTACAAACA GATCTAGCGAAAATTTACCATTGTCATGTTTACATGTTTAAACATCCCTCTTTG CCAGTGTTAATTACCAGAATACAGCTATTTGACAGCAATAATTTATTTTTGAGT ACGCCTAATATAGGGAGTATAAATAAAGAGTCTCTTTACAACAAGTTAGACAA ATTCCAAGGGAAACCATTAATATCAAGGAGtCCATATTACGTTGCATTCCCGC TAAATTCTCCAATAATTTTCCATTCAGTTGACAgtGATATTTATGCTAGACTAGT TTTGCAGAGCATTAGTCGAACGATATCAGAAAGAGAAACAATCATCTTCAAA CCGGTACAGtcAATACCCGTGtcATCAATACATAATATAATGACTTTATTAGGAC CTTCGAGATTTGCGGAGTCTATGGGCCCATGGGAATGTTATGCTAGTGCTAA CTTTGAACGGTCACCACTACACGACTACAAAAAGCACCAAGGCCTTACTGGT AAAAAGGTAATGGTTAGAGAATTTGATGATTCTTTCTTAAATGACGATGAGAA TTTTTACGGAAAAGAAGAACCTGAAATTAGACGACTAAGGTTAGAGAAAAAC ATGATCAAATTCAAAGGTTCAGCAAACGGCGTAATGGATCAGAAATATAATG ATTTGAAAGAATTTAACGAACATGTCCACAATATTAGGAATGGAAAGAAAAAT GAAGATTCTGGTGAGCCAGTGTACATTTCTAGATACAGTTCCTTAGTTCCTAT TGAAtcAGTTGGATTCACTTTAAAAAACGAAATAAACAGTAGAATAATTACCAT AtcATTGAgtTTTAATGGTAATGATATTTTTGGGGGATTACACGAACTATGCG - 3' | gBlock to mutagenize *CHL4* at 13 K/R residues that contact DNA (K/R- >S) |

| | | |
|---|---|---|
| SB7872 | 5' - AGTACGAGGCCAAACTCTCGAAAAGGATATTACGAGATGCGGCCGCTCTAGAACTAGTGG- 3' | 5' primer to tag *SCM3* with mCherry with pSB3167 |
| SB3878 | 5' - biotin-GGTTCTGGTGGTTCTGGTGGTTCTGGTGAATTCAAACAACCGCCGGCTTCCACCA - 3' | 250 bp single-tether CEN3 or double-tether or 250bp CEN3$^{mut}$ templates |
| SB3879 | 5' - biotin-GGTTCTGGTGGTTCTGGTGGTTCTGGTGGAATTCATTGTTTGTGCACTTGCCTGCA - 3' | 750 bp CEN3, CEN3$^{mut}$, CEN9, CEN3-CDEII-L1, CEN3-CDEII-L5, CEN3-CDEII-H1, CEN3-CDEII-H14, or CEN3-CDEII-$\alpha$-sat templates |
| SB3880 | 5' - ATCAGCGCCAAACAATATGGAA - 3' | 250 bp single-tether CEN3 or 250bp CEN3$^{mut}$ templates |
| SB5699 | 5' - biotin-GGTTCTGGTGGTTCTGGTGGTTCTGGTGAATTCATGAAATCAAAATTAAACATTTT - 3' | 750 bp CEN7 template |
| SB7054 | 5' - biotin-GGTTCTGGTGGTTCTGGTGGTTCTGGTGAATTCCATTAATGCAGCTGGCACGAC - 3' | 750 bp Widom-601 and Widom-601 hybrid templates |

| | | |
|---|---|---|
| SB7842 | 5' - ATTO647-CTCTAAAGAAATTGAAAACTTG - 3' | 750 bp CEN7 template |
| SB7870 | 5' - ATTO647-ATGGTGTTTATGCAAAGAAACCA - 3' | 750 bp CEN3, CEN3$^{mut}$, CEN9, CEN3-CDEII-L1, CEN3-CDEII-L5, CEN3-CDEII-H1, CEN3-CDEII-H14, or CEN3-CDEII-α-sat templates |
| SB7871 | 5' - ATTO647-AGAAAATACCGCATCAGGCGC - 3' | 750 bp Widom-601 and Widom-601 hybrid templates |
